# Predicting alcohol-related memory problems in older adults: A machine learning study with multi-domain features

**DOI:** 10.1101/2022.12.30.522330

**Authors:** Chella Kamarajan, Ashwini K. Pandey, David B. Chorlian, Jacquelyn L. Meyers, Sivan Kinreich, Gayathri Pandey, Stacey Subbie-Saenz de Viteri, Jian Zhang, Weipeng Kuang, Peter B. Barr, Fazil Aliev, Andrey P. Anokhin, Martin H. Plawecki, Samuel Kuperman, Laura Almasy, Alison Merikangas, Sarah J. Brislin, Lance Bauer, Victor Hesselbrock, Grace Chan, John Kramer, Dongbing Lai, Sarah Hartz, Laura J. Bierut, Vivia V. McCutcheon, Kathleen K. Bucholz, Danielle M. Dick, Marc A. Schuckit, Howard J. Edenberg, Bernice Porjesz

## Abstract

Memory problems are common among older adults with a history of alcohol use disorder (AUD). Employing a machine learning framework, the current study investigates the use of multi-domain features to classify individuals with and without alcohol-induced memory problems. A group of 94 individuals (ages 50-81 years) with alcohol-induced memory problems (*Memory* group) were compared with a matched *Control* group who did not have memory problems. The Random Forests model identified specific features from each domain that contributed to the classification of Memory vs. Control group (AUC=88.29%). Specifically, individuals from the Memory group manifested a predominant pattern of hyperconnectivity across the default mode network regions except some connections involving anterior cingulate cortex which were predominantly hypoconnected. Other significant contributing features were (i) polygenic risk scores for AUD, (ii) alcohol consumption and related health consequences during the past 5 years, such as health problems, past negative experiences, withdrawal symptoms, and the largest number of drinks in a day during the past 12 months, and (iii) elevated neuroticism and increased harm avoidance, and fewer positive “uplift” life events. At the neural systems level, hyperconnectivity across the default mode network regions, including the connections across the hippocampal hub regions, in individuals with memory problems may indicate dysregulation in neural information processing. Overall, the study outlines the importance of utilizing multidomain features, consisting of resting-state brain connectivity collected ∼18 years ago, together with personality, life experiences, polygenic risk, and alcohol consumption and related consequences, to predict alcohol-related memory problems that arise in later life.

## 1. Introduction

Alcohol use disorder (AUD) is a chronic, relapsing disorder [1, 2] with a range of neurocognitive anomalies, including memory deficits [3]. Memory impairments due to heavy drinking, among other cognitive impairments, have been widely reported [4, 5], and may interfere with social and occupational performance [6, 7]. Since the etiology of AUD and related memory problems involves multiple domains, including the combination of neurocognitive, personality, behavioral, and genomic factors [8–10], a better understanding of these potential predictors may aid in prevention and treatment strategies.

Brain oscillations representing electrical signals of neural activity, as recorded by electroencephalogram (EEG), index specific circuit-level mechanisms during cognitive processing [11]. Oscillatory signals in different EEG frequency bands representing communications between specific brain regions underlie memory processes, including encoding, consolidation, storage, and retrieval processes [12, 13]. Studies have indicated that memory processes are supported by oscillatory dynamics and communication across the hippocampus, entorhinal cortex and other cortical regions [13–15]. Both human and animal studies have implicated the theta band, generated within the hippocampus and also prevalent in the cerebral cortex, as the major frequencies associated with various memory processes [16, 17]. Hippocampal theta rhythm is also involved in communication with other higher frequencies (e.g., beta and gamma oscillations) through various coupling mechanisms, including neural synchrony during sensory and cognitive processing [18–21].

Recent studies have used source localization methods, such as the exact low-resolution brain electromagnetic tomography (eLORETA) [22] to compute *functional connectivity* a measure of temporal synchrony or correlation between signals of two or more spatially separated brain regions representing functional integration between these areas [cf. 23]. These studies have used *lagged connectivity* [24] to overcome volume conduction artifacts [23, 25]. While the eLORETA-based functional connectivity method has been utilized to study cognitive functioning in neuropsychiatric disorders [23,26–29], very few studies have utilized these approaches to investigate AUD [30] and none of these studies have examined alcohol-induced neurocognitive outcomes, such as memory. Since the default mode network supports memory functions [31–34], we will employ functional connectivity across the default mode network regions to examine alcohol-related neurocognitive outcomes such as memory impairment to include features from multiple domains, including polygenic risk scores (PRS) [35, 36] and personality dimensions [36–41]. Therefore, the goal of the present study was to identify a set of multi-domain factors that can differentiate individuals with alcohol-related memory impairments from those without, using (i) resting EEG-based functional connectivity measures of default mode network as derived from eLORETA, (ii) PRS related to alcohol outcomes, (iii) personality and life experience measures derived from established questionnaires, and (iv) measures of alcohol consumption and associated health consequences from the recent follow-up interview. Identifying specific default mode network functional connections underlying alcohol-induced memory problems may be useful for early preventive measures and for brain-based treatment strategies such as neuromodulation therapies for addiction [42] and memory/cognitive impairment or decline [43]. Similarly, other domains, including PRS, behavioral, personality, and clinical features, may have implications for prevention and treatment of alcohol-induced memory problems (e.g., cognitive-behavior therapy, brain stimulation, cognitive remediation, etc.).

## 2. Material and Methods

### 2.1. Sample

The sample for the present study was drawn from a recent follow-up assessment study [44, 45] of participants from the Collaborative Study on Genetics of Alcoholism (COGA) [46–48]. Participants aged 50 or older who met lifetime criteria for alcohol dependence, as assessed with the Semi-Structured Assessment for the Genetics of Alcohol (SSAGA) [49, 50], were drawn from data collected at six COGA sites. Details on screening and selection of participants for the current study are described the supplemental material [see *Section 1.1. Sample Description* and *Fig. S1* in the *Supplementary Material*]. The *Memory* and *Control* groups were also matched for age at assessments, sex, self-reported race, genetic ancestry, and the following alcohol use patterns assessed by their last SSAGA interview conducted ∼18 years prior to the recent telephone interview (see **Table 1**): (i) continued high-risk drinking (men with 5+ drinks/day or than 5 drinks/day for men and 4 drinks/day for women) without meeting criteria for AUD diagnosis (N=9/group), and (iii) abstinence from drinking (N=17/group).

**Table 1:** Demographic characteristics, AUD remission status during the latest SSAGA interview before the follow up telephone interview, and details of alcohol consumption from the recent telephone interview for the EEG functional connectivity analysis.

| Variable | Measure / Category | Parameter | Study Group |  |
| --- | --- | --- | --- | --- |
|  |  |  | Memory (N=94) | Control (N=94) |
| Age during assessment | EEG* | Min–Max | 29.21–60.71 | 28.17–62.19 |
|  |  | Mean (SD) | 39.42 (6.18) | 40.11 (6.74) |
|  | Follow-up Interview | Min–Max | 50.55–81.86 | 50.34–81.49 |
|  |  | Mean (SD) | 57.84 (5.77) | 58.75 (6.07) |
| Sex | Male | N (%) | 52 (55.30) | 52 (55.30) |
|  | Female | N (%) | 42 (44.70) | 42 (44.70) |
| Self-reported race | White | N (%) | 67 (71.30) | 67 (71.30) |
|  | Black | N (%) | 24 (25.50) | 24 (25.5) |
|  | Other | N (%) | 3 (3.20) | 3 (3.20) |
| Genetic ancestry | European | N (%) | 63 (50.40) | 62 (49.60) |
|  | African | N (%) | 23 (47.92) | 25 (52.08) |
|  | Other | N (%) | 8 (53.33) | 7 (46.67) |
| Alcohol use pattern during the latest SSAGA interview* | AUD diagnosis | N (%) | 68 (72.30) | 68 (72.30) |
|  | Low Risk Drinking | N (%) | 9 (9.60) | 9 (9.60) |
|  | Abstinence | N (%) | 17 (18.10) | 17 (18.10) |
| Time lag** | Years | Mean (SD) | 18.42 (3.84) | 18.63 (3.90) |

\*The latest SSAGA interviews were also closer in time to the EEG recording used for the current study. Note that the SSAGA interview is longer and more comprehensive than the recent follow-up phone interview.
\*\* Time lag (years) between the latest past (baseline) assessments (EEG, SSAGA, and clinical/personality) and the recent follow-up telephone interview.

### 2.2. Recent Telephone Interview

The recent follow-up telephone interview (10-20 minutes) was designed to collect information regarding participants’ alcohol use and current social and health status using a 31-items questionnaire [45] administered via the REDCap system [51, 52]. Details about this interview items are available in *Section 1.2 of the Supplementary Material*. Three items that elicited self-reported alcohol-related memory problems have been listed in **Table 2**. Memory impairment was coded if the participant endorsed at least two of the three items (**Table 2**): the first item and either the second or third item.

**Table 2:** Items related to memory problems in the follow-up interview questionnaire.

| Domain | Question | Memory-related response* |
| --- | --- | --- |
| **Alcohol-related memory problems** | Compared to most people your age, is your memory currently better, about the same, or worse than theirs? | ○ Worse |
|  | **There are several other health problems that can result from heavy drinking. In the last 5 years did drinking. (Check all that apply) | ○ Impair your memory even when you were not drinking (not including blackouts)? |
|  | **There are several other health problems that can result from heavy drinking. In the last 10 years did drinking. (Check all that apply) | ○ Impair your memory even when you were not drinking (not including blackouts)? |

\* Response option related to memory problems.
\*\* These items are the same for the categories eliciting alcohol use during the past 5 years and past 10 years.

### 2.3. EEG Data Acquisition and Preprocessing

Details of assessments and EEG recording in COGA, which is identical at all sites, can be found in our previous reports [46,53,54]. The EEG session that was closest to the latest SSAGA interview was used for this study. Detailed descriptions of EEG data acquisition and preprocessing steps are available in *Section 1.3* of the *Supplementary Material*.

### 2.4. EEG Functional Connectivity Analysis using eLORETA

EEG functional connectivity was computed using the eLORETA software [22, 55], a validated tool for localizing the electrical activity in the brain. Detailed descriptions of EEG functional connectivity analysis using eLORETA are available in the *Section 1.4* of the *Supplementary Material*.

### 2.5. Functional Connectivity Across the Default Mode Network

The default mode network regions analyzed in the study are posterior cingulate cortex (PCC), anterior cingulate cortex, inferior parietal cortex, prefrontal cortex, lateral temporal cortex, and hippocampal formation [see **Table 3** below and **Fig. S2** in the *Supplementary Material*], in line with the functional connectivity studies of both fMRI and EEG [28,56,57] and our previous work on default mode network [58, 59].

**Table 3.** Regions of interest (ROI), region code/abbreviation, Brodmann area (BA) and the MNI coordinates for the default mode network are listed.

| ROI | Region Name | Region Code | BA | MNI (X) | MNI (Y) | MNI (Z) |
| --- | --- | --- | --- | --- | --- | --- |
| 1 | Left posterior cingulate cortex | L.PCC | 23 | -10 | -45 | 25 |
| 2 | Right posterior cingulate cortex | R.PCC | 23 | 10 | -45 | 25 |
| 3 | Left anterior cingulate cortex | L.ACC | 32 | -10 | 45 | 10 |
| 4 | Right anterior cingulate cortex | R.ACC | 32 | 10 | 45 | 10 |
| 5 | Left inferior parietal lobule | L.IPL | 40 | -55 | -55 | 20 |
| 6 | Right inferior parietal lobule | R.IPL | 40 | 55 | -55 | 20 |
| 7 | Left prefrontal cortex | L.PFC | 46 | -45 | 25 | 25 |
| 8 | Right prefrontal cortex | R.PFC | 46 | 45 | 25 | 25 |
| 9 | Left lateral temporal cortex | L.LTC | 21 | -55 | -15 | -20 |
| 10 | Right lateral temporal cortex | R.LTC | 21 | 55 | -15 | -20 |
| 11 | Left parahippocampal gyrus | L.PHG | 36 | -25 | -30 | -20 |
| 12 | Right parahippocampal gyrus | R.PHG | 36 | 25 | -30 | -20 |

### 2.6. Assessment of Temperament, Personality, and Alcohol Experience

The temperament, personality, and life experiences data included scores from seven questionnaires and their subscales, and scores included for the current study are described in *Section 1.6* of the *Supplementary Material.* These data were collected during the previous interviews (∼18 years ago) at/around the same time as the SSAGA assessment.

### 2.7. Genomic Data and Polygenic Risk Scores (PRS)

Genotyping, imputation, and quality control of COGA genomic data have been described previously [48] and in the *Section 1.7* of the *Supplementary Material*. The publicly available Genome-wide Association Studies (GWAS) for alcohol use phenotypes, derived from studies including both individuals of European ancestry (EA) and African ancestry (AA), that were used in PRS calculations in this study are listed in **Table 4**.

**Table 4.** List of Polygenic Risk Scores (PRS) datasets from recently published GWAS

| Phenotype | Discovery<br>Sample/Consortium | Sample Size |  |
| --- | --- | --- | --- |
|  |  | EA | AA |
| AUD diagnosis (ICD-9/ICD-10) | MVP [60] | 202,004 | 56,648 |
| AUDIT-C symptoms | MVP [60] | 200,680 | 56,495 |
| Max alcohol intake | MVP [61] | 126,936 | 17,029 |
| Alcohol Dependence (DSM-IV) | PGC [62] | 46,568 | 6,280 |

We created PRS using PRS-CSx [63–67], which is a recent, validated method for cross-ancestry polygenic prediction [68]. The PRS-CSx computation method is detailed elsewhere (https://github.com/getian107/PRScsx) and also briefly described in the *Section 1.7* of the *Supplementary Material*.

### 2.8. Feature selection of EEG functional connectivity variables

In keeping with recent machine learning approaches, including our previous study [69], we used a two-stage approach consisting of feature selection followed by a predictive algorithm using selected sets of variables [70–74]. A detailed description of this method is available in *Section 1.8* of the *Supplementary Material*.

### 2.9. Random Forests classification model and parameters

The Random Forests classification analysis was performed using R-packages “randomForest” [75], “caret” [76], and “randomForestExplainer” [77] to classify *Memory* vs. *Control* group using multi-domain predictors. The details of these predictors, which include 29 functional connectivity, 27 personality and life experience, 12 alcohol outcomes, and 4 PRS variables, are listed in the *Materials and Methods* section of the *Supplementary Material*. The random forests model, as implemented in the current study, has been detailed in *Section 1.9* of the *Supplementary Material*.

## 3. Results

### 3.1. Feature Selection of EEG functional connectivity variables

The input data for the feature selection included a total of 330 EEG functional connectivity variables consisting of 66 connectivity features for each of five frequency bands. The model identified a total of 29 functional connectivity variables from multiple frequency bands connecting across the twelve default mode network seeds (Refer **Table 3** in *Methods* section and **Fig. S2** in *Supplementary Material*). These connections included Delta – 12 connections, Theta – 6 connections, Alpha – 4 connections, Beta – 5 connections, and Gamma – 2 connections. The 10-fold cross-validation for the λ1se threshold included all the 29 selected features, which were included in the subsequent implementation of the Random Forests classification model. The classification performance (to differentiate individuals with memory problems from those without) of the selected features as indicated by the area under the ROC curve (AUC) was 88.48%.

### 3.2. Random Forests Classification Accuracy

The overall prediction accuracy of the Random Forests model to classify *Memory* and *Control* group using functional connectivity, PRS, behavioral and clinical predictors, as estimated by the AUC, was 88.29%. The 72 predictors inputted in the model include 29 functional connectivity, 27 personality and life experience, 12 alcohol outcomes, and 4 PRS variables (see *Materials and Methods* section of the *Supplementary Material*). Additional details about the classification accuracy are available in *Section 2.2*. of the *Supplementary Material*.

### 3.3. Top Significant Features Contributed to the Classification

Out of the 72 input variables of the Random Forest model (see *Materials and Methods* section of the *Supplementary Material* for details), 29 significant features that contributed to classifying *Memory* group from those from the *Control* group were identified: 21 default mode network connections, 4 alcohol-related items, 3 personality and life experience factors, and 1 PRS (**Table 5**).

**Table 5.** Random Forest importance parameters and direction of significance for the top significant variables (p < 0.05) are shown. The variables are sorted based on Gini decrease. Details of these features are available in *Materials and Methods* section of the *Supplementary Material*.

| Feature | Measure / Source | Gini Decrease | Accuracy Decrease | # Trees | # Nodes | Times a Root | Min. Depth | P value | Direction |
| --- | --- | --- | --- | --- | --- | --- | --- | --- | --- |
| AlcHlthProb5yrs | FU Interview | 7.7281 | 0.0449 | 545 | 610 | 111 | 2.3303 | 8.26E-47 | MEM > CTL |
| AlcWthSx5yrs | FU Interview | 4.8291 | 0.0196 | 430 | 459 | 109 | 3.8230 | 4.09E-13 | MEM > CTL |
| AlcExp5yrs | FU Interview | 4.8134 | 0.0176 | 417 | 468 | 95 | 4.0144 | 1.42E-14 | MEM > CTL |
| Drk24Hr | FU Interview | 2.7318 | 0.0097 | 385 | 440 | 70 | 5.0280 | 2.75E-10 | MEM > CTL |
| *NEO_N | Questionnaire | 1.9701 | 0.0029 | 334 | 382 | 47 | 5.6475 | 6.84E-04 | MEM > CTL |
| FC_Ga_2_10 | R.PCC–R.LTC | 1.9574 | 0.0019 | 402 | 486 | 5 | 5.5047 | 1.02E-17 | MEM > CTL |
| FC_Th_2_11 | R.PCC–L.PHG | 1.8902 | 0.0020 | 377 | 463 | 11 | 5.7415 | 9.38E-14 | MEM > CTL |
| FC_Be_1_4 | L.PCC–R.ACC | 1.8699 | 0.0030 | 378 | 463 | 6 | 5.8232 | 9.38E-14 | CTL > MEM |
| FC_Th_2_5 | R.PCC–L.IPL | 1.7564 | 0.0039 | 356 | 424 | 16 | 5.8446 | 3.53E-08 | MEM > CTL |
| FC_Th_9_11 | L.LTC–L.PHG | 1.7206 | 0.0010 | 362 | 437 | 17 | 5.8282 | 7.15E-10 | MEM > CTL |
| FC_De_1_5 | L.PCC–L.IPL | 1.6655 | 0.0011 | 346 | 412 | 12 | 6.0057 | 9.11E-07 | MEM > CTL |
| *TPQ_HA | Questionnaire | 1.6312 | 0.0026 | 318 | 363 | 37 | 6.1333 | 1.44E-02 | MEM > CTL |
| FC_Al_2_5 | R.PCC–L.IPL | 1.6034 | 0.0013 | 376 | 455 | 9 | 5.9314 | 1.72E-12 | MEM > CTL |
| FC_De_2_5 | R.PCC–L.IPL | 1.5614 | 0.0004 | 366 | 437 | 18 | 5.8339 | 7.15E-10 | MEM > CTL |
| FC_De_1_6 | L.PCC–R.IPL | 1.5384 | 0.0009 | 310 | 383 | 27 | 6.2101 | 5.68E-04 | MEM > CTL |
| FC_Be_4_9 | R.ACC–L.LTC | 1.4901 | 0.0009 | 344 | 402 | 12 | 6.2038 | 1.05E-05 | CTL > MEM |
| FC_Ga_4_12 | R.ACC–R.PHG | 1.4605 | 0.0016 | 376 | 451 | 3 | 5.6709 | 6.99E-12 | CTL > MEM |
| FC_De_7_11 | L.PFC–L.PHG | 1.4543 | 0.0019 | 342 | 407 | 13 | 6.1891 | 3.19E-06 | MEM > CTL |
| *DHU_UPL | Questionnaire | 1.4497 | 0.0021 | 315 | 368 | 15 | 6.4736 | 7.06E-03 | CTL > MEM |
| FC_Th_4_10 | R.ACC–R.LTC | 1.4211 | 0.0006 | 345 | 422 | 8 | 6.2084 | 6.21E-08 | CTL > MEM |
| FC_De_8_12 | R.PFC–R.PHG | 1.3844 | 0.0010 | 333 | 394 | 15 | 6.0851 | 6.29E-05 | MEM > CTL |
| FC_Al_2_11 | R.PCC–L.PHG | 1.3805 | 0.0006 | 360 | 443 | 3 | 6.2337 | 1.04E-10 | MEM > CTL |
| PRS_MVP_AUD | PRS | 1.2987 | 0.0002 | 363 | 432 | 1 | 6.2696 | 3.35E-09 | CTL > MEM |
| FC_De_5_6 | L.IPL–R.IPL | 1.2964 | 0.0009 | 320 | 378 | 11 | 6.4012 | 1.40E-03 | MEM > CTL |
| FC_De_6_11 | R.IPL–L.PHG | 1.2959 | -0.0001 | 317 | 381 | 10 | 6.3433 | 8.21E-04 | MEM > CTL |
| FC_Th_4_6 | R.ACC–R.IPL | 1.2955 | 0.0002 | 342 | 404 | 2 | 6.3120 | 6.59E-06 | CTL > MEM |
| FC_De_2_12 | R.PCC–R.PHG | 1.2581 | 0.0007 | 319 | 380 | 9 | 6.4407 | 9.83E-04 | MEM > CTL |
| FC_De_4_8 | R.ACC–R.PFC | 1.1741 | 0.0015 | 315 | 364 | 6 | 6.5837 | 1.26E-02 | MEM > CTL |
| FC_De_3_7 | L.ACC–L.PFC | 1.1278 | 0.0000 | 319 | 391 | 6 | 6.7618 | 1.18E-04 | CTL > MEM |

**Abbreviations:** FC–Functional Connectivity; De–Delta; Th–Theta; Al–Alpha; Be–Beta; Ga–Gamma; Numbers in functional connectivity variables: 1-12 of the default mode network; AlcHlthProb5yrs–Alcohol-related health problems in the past 5 years; AlcWthSx5yrs–Alcohol withdrawal symptoms in the past 5 years; AlcExp5y–Alcohol related negative experiences (symptoms) related to alcohol consumption in the past 5 years; Drk24Hr–The largest number of drinks in 24 hours during the past 12 months; PRS\_MVP\_AUD–PRS derived from the MVP GWAS of AUD; \*Measures from personality and life experience questionnaires: TPQ\_HA–Harm avoidance assessed by TPQ questionnaire; DHU\_UPL–Uplift assessed by DHU questionnaire; and NEO\_N–Neuroticism assessed by NEO questionnaire [See **Table 3** in Methods and **Fig. S2** in the *Supplementary Material* for the details of the ROIs of the default mode network]. MEM–Memory group; CTL–Control group

The multi-way importance plot [**Fig. 1**] displays all significant variables (labeled and marked with black circles) that contributed to the classification of the *Memory* group from the *Control* subjects and ranked based on the importance for classification as derived from Gini decrease, number of trees, and p-value. A chart showing distribution of minimal depth in classification against number of decision trees [see **Fig. S4** in the *Supplementary Material*]. While both multi-way importance plot and distribution plot can be created for any set of random forest parameters, the importance ranking for the features is likely to be similar owing to high correlations among these parameters (see **Fig. S5** in *Supplementary Material*).

**Fig. 1.**
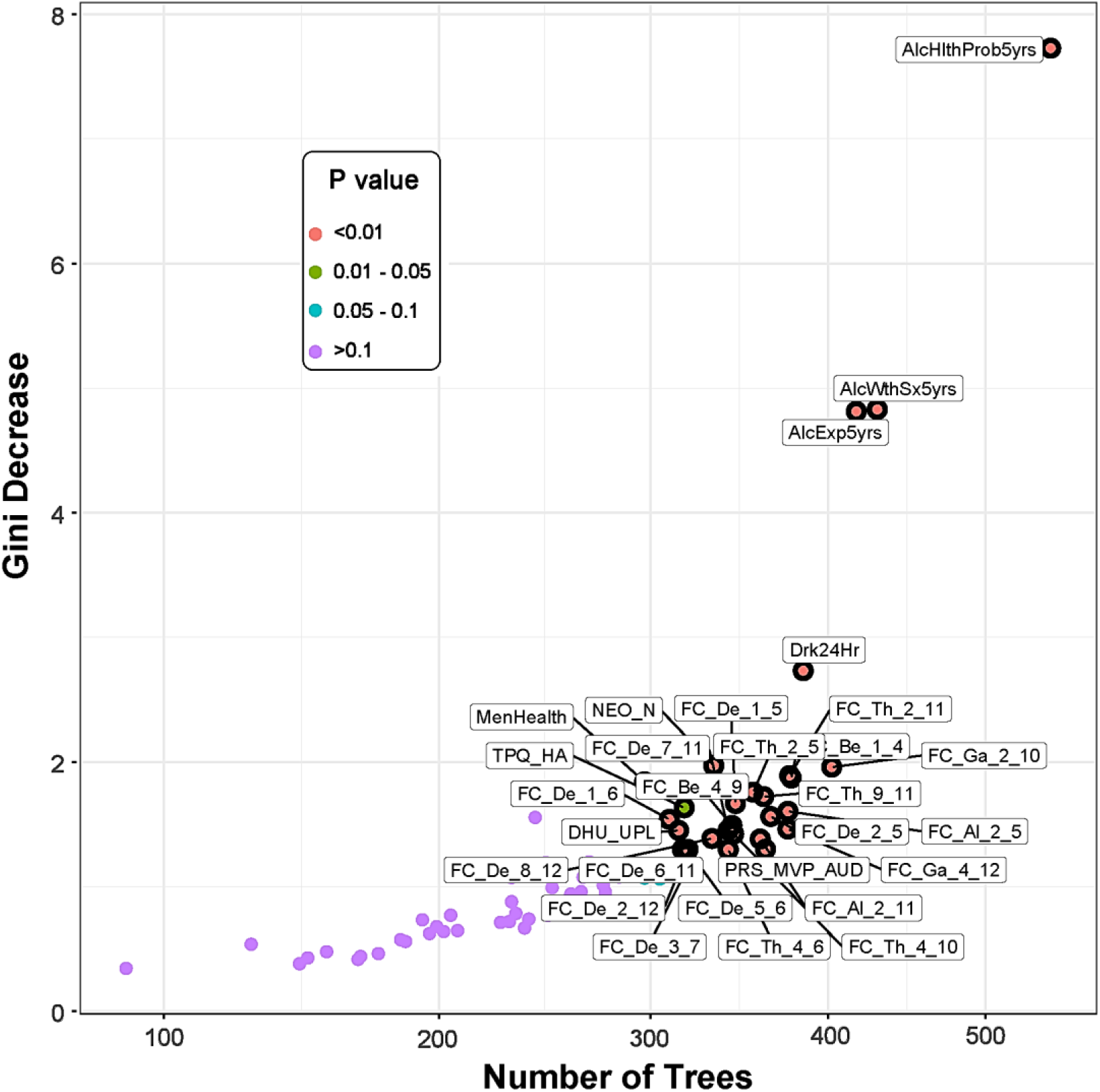
**The multi-way importance plot showing the top significant variables (labeled and marked with black circles) that contributed to the classification of the *Memory* group from the *Control* subjects based on the measures Gini decrease, number of trees, and p-value:** Features related to alcohol-related clinical/health outcomes stood top in the importance list, followed by functional connectivity, personality, and PRS measures. Note that the variables that were not significant (purple dots) are not highlighted. [See footnote of **Table 5** for the list of abbreviations for the measures shown here].

#### 3.3.1. EEG Source Functional Connectivity of the Default Mode Network

Significant default mode network connections which contributed to the Random Forest classification of the *Memory* group from *Control* individuals have been illustrated in **Fig. 2**. *Memory* group showed a predominant pattern of hyperconnectivity across the default mode network regions, primarily contributed by delta band (10 connections) followed by theta band (5 connections) band, along with fewer hypoconnectivity (1 in delta band and 2 in theta band). Other significant functional connectivity features specific to each frequency band are (i) 9 hyperconnected paths and 1 hypoconnected path in delta band, (ii) 3 hyperconnected and 2 hypoconnected paths in theta band, (iii) 2 hyperconnected paths with no hypoconnected paths in alpha band, (iv) 2 hypoconnected paths with no hyperconnected paths in beta band, and (v) 1 hyperconnected path and 1 hypoconnected path gamma band (**Fig. 2, Panels A-E**). Number of significant connections from each ROI node (in descending order) was as follows: R.PCC = 7; R.ACC = 6; L.PHG = 5; L.IPL = 5; R.IPL = 4; L.PCC = 3; R.PHG = 3; L.PFC = 2; R.PFC = 2; L.LTC = 2; R.LTC = 2; L.ACC = 1. The number of significant connections for the ROIs involving both hemispheres (in ascending order) was: PCC = 10; IPL = 9; PHG = 8; ACC = 7; PFC = 4; LTC = 4. Individuals from *Memory* group showed predominant hyperconnectivity between hippocampal region (PHG) and other default mode network regions involving multiple frequencies except beta band compared with the *Control* group (**Fig. 2, Panel F**). Only a single hippocampal connection (R.PHG–R.ACC) of the gamma band oscillation was hypoconnected in the *Memory* group.

**Fig. 2:**
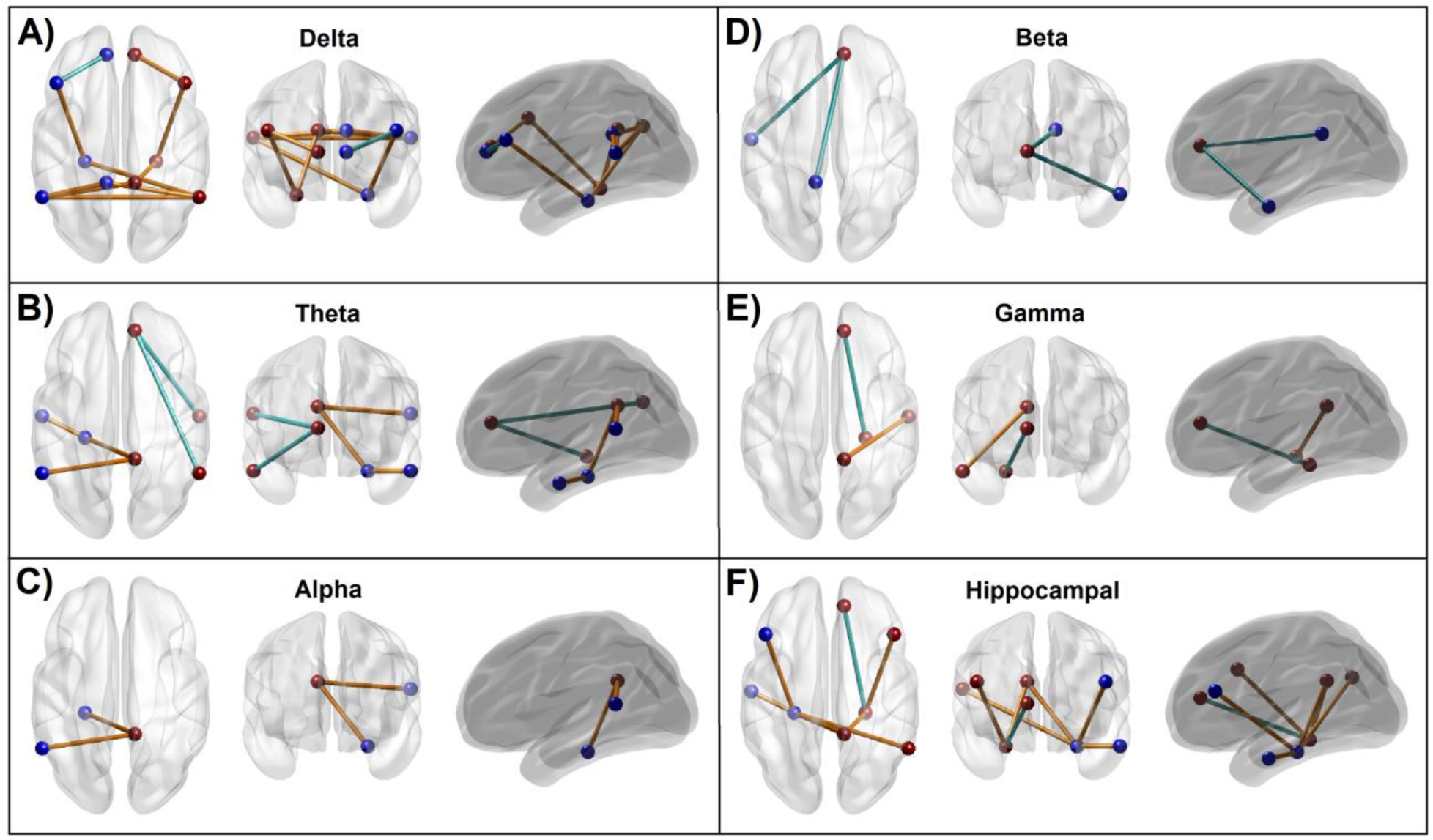
***Panels A-E:*** Significant default mode network connections within each frequency band, which contributed to the Random Forest classification of *Memory* group from *Control* individuals. The blue and brown beads represent ROIs of the left and right hemisphere, respectively, while the blue and brown lines represent hypoconnectivity and hyperconnectivity, respectively, in the *Memory* group. ***Panel F:*** Significant hippocampal connections that contributed to the *Memory* vs. *Control* classification. Seven of the eight hippocampal connections showed hyperconnectivity in the *Memory* group. Note that all hypoconnected networks involved an anterior cingulate node. Refer to *Fig. S2* in the *Supplementary Material* for the ROI locations and anatomical views/axes.

#### 3.3.2. Recent Alcohol Consumption and Health Outcomes

Significant alcohol-related health outcome variables that contributed to classifying *Memory* individuals from the *Control* subjects included (i) alcohol-related health problems in the past 5 years (*Memory*_mean_=0.77; *Control*_mean_=0.01), (ii) alcohol withdrawal symptoms in the past 5 years (*Memory*_mean_=1.20; *Control*_mean_=0.11), (iii) negative experiences related to alcohol consumption in the past 5 years (*Memory*_mean_=2.65; *Control*_mean_=0.78), and (iv) the largest number of drinks within 24 hours during the past 12 months (*Memory*_mean_=13.64; *Control*_mean_=6.00). Interestingly, the features concerning alcohol-related outcomes over the past 10 years, physical health outcomes, other drinking patterns, and demographic variables were not significant.

#### 3.3.3. Measures of Personality, Behavior, and Life Experiences

Out of 27 variables of personality and behavioral features, only the following three variables significantly contributed to the *Memory* vs. *Control* classification: (i) Harm avoidance representing internalizing traits and negative mood states as assessed by TPQ (*Memory*_mean_=16.16; *Control*_mean_=12.61), (ii) Uplift experience indicating “feel good” aspects as assessed by DHU (*Memory*_mean_=51.25; *Control*_mean_=58.99), and (iii) Neuroticism represented by dysregulated emotions and maladjusted behaviors as assessed by NEO (*Memory*_mean_=59.00; *Control*_mean_=52.11), and higher scores mean more neurotic traits.

#### 3.3.4. Polygenic Risk Scores

PRS for the AUD diagnosis (based on the ICD codes) created using GWAS data from the MVP [60] was a significant contributor to the classification of *Memory* vs. *Control* group (*Memory*_mean_=8.25 × 10^-7^ and *Control*_mean_=7.87 × 10^-7^). PRSs for the other phenotypes, i.e., AUDIT-C scores from the GWAS of MVP dataset [60], Maximum habitual alcohol intake from the GWAS of MVP dataset [61], and DSM-IV alcohol dependence diagnosis from the GWAS of PGC dataset [62], were not significant contributors in the classification.

### 3.4. Correlations across Significant Predictors

Exploratory (descriptive) analysis of correlations among the top significant variables is shown in **Fig. 3**. As shown in the correction matrix, there were significant positive correlations relationships among the functional connectivity variables within and between different frequency bands. Overall, most of the low-frequency connections in the delta and theta frequencies were highly correlated with one another. Specifically, those connections that shared a common node showed much higher correlations with each other than with other connections, regardless of their frequency band. Beta band connections had significant positive correlations between themselves as well as with low-frequency connections, especially that of theta band connections. However, alpha and gamma band connections showed significant correlations only within the frequency but not across the frequencies. Highly significant positive correlations were observed among the alcohol-related health consequences. Among the personality factors, there was a significant positive correlation between neuroticism and harm avoidance. However, no significant correlations were observed across the domains (e.g., functional connectivity vs. personality, or functional connectivity vs. alcohol-related features).

**Fig. 3:**
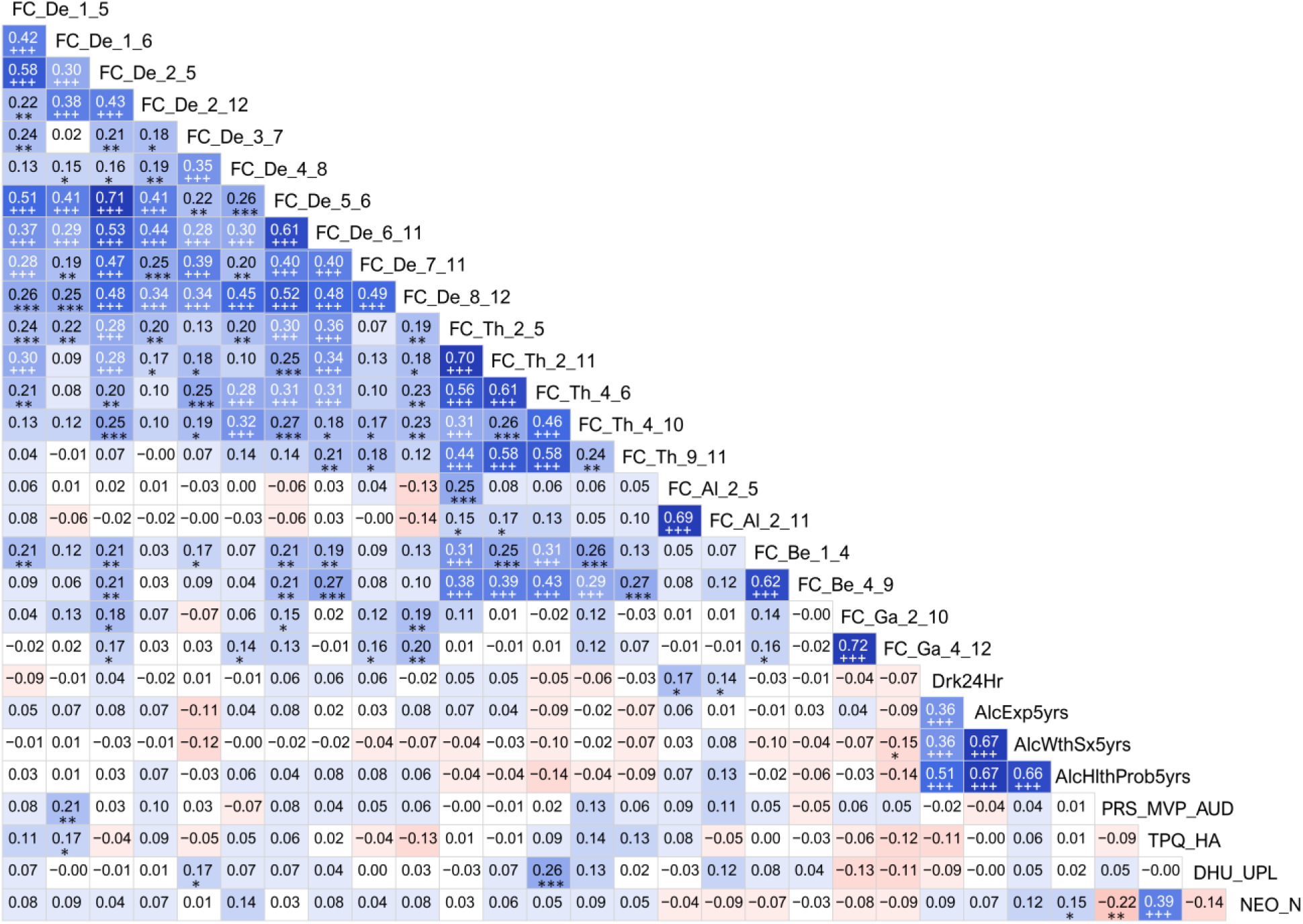
Correlation matrix showing associations among the top significant variables. Values of the cells in red/pink shades represent negative r-values, and those in blue/cyan shades indicate positive r-values between variables that correspond to the vertical and horizontal axis. Darker color represents a higher magnitude of r-values. Significant correlations (before Bonferroni correction) have been marked with asterisks in black font [*p < 0.05; **p < 0.01; and ***p < 0.001], and those survived Bonferroni correction have been marked with a triple plus sign (+++) in white font. For the abbreviations in the variable labels, see the footnote of **Table 5**.

## 4. Discussion

The current study suggests that alcohol-related memory problems can be predicted using a multi-domain set of features from neural, behavioral, genomic, and alcohol-related measures in a machine learning framework. It was found that the *Memory* group showed a predominant pattern of hyperconnectivity across the default mode network regions, including the hippocampal subnetworks, while showing hypoconnected anterior cingulate cortex subnetworks based on the EEG recorded about 18 years ago. Features from other domains that significantly contributed to the classification were (i) higher counts of alcohol-related consequences during the past 5 years, such as health problems, other alcohol-related adverse past negative experiences, withdrawal symptoms, and higher max number of drinks (the largest number of drinks per day), (iii) personality factors such as high neuroticism, high harm avoidance, and low positive/uplift experience, and (iv) high genetic liability, as reflected in variations in PRS for AUD across the *Memory* and *Control* groups. It should also be noted that the classification accuracy was better for the *Control* individuals (85/94 = 90.43%) than for the *Memory* group (68/94 = 72.34%). Although the reasons could be many, we speculate that the *Memory* group may have high variability in their clinical presentations and/or neurocognitive functioning.

### 4.1. Altered Functional Connectivity in the Memory group

Findings of resting-state EEG connectivity showed that those with alcohol-related memory problems, relative to matched controls, showed (i) a predominant pattern of hyperconnectivity of low-frequency (delta and theta) oscillations across most of the default mode network cortical regions, (ii) hyperconnected hippocampal sub-networks in multiple frequency bands, and (iii) hypoconnectivity in subnetworks involving anterior cingulate cortex hub regions. In general, alterations in brain networks (in both low and high frequencies) due to alcohol-induced memory deficits could be interpreted as compromised memory engrams and changes in neural plasticity during encoding and recall processes. The neural basis of memory processes was first theorized by Richard Semon’s *engram theory* [78] and Donald Hebb’s *synaptic plasticity theory* [79] and here is a vast literature spanning several decades on memory functions. The connectivity differences observed between *Memory* and *Control* groups are discussed below in light of findings from the literature as well as our previous studies.

#### 4.1.1. Predominant hyperconnectivity of low-frequency oscillations in the Memory group

The finding that individuals with alcohol-induced memory problems during their recent interview (i.e., *Memory* group) manifested a predominant pattern of hyperconnectivity across the default mode network nodes in their resting state EEG [**Fig. 2**] may indicate aberrations in neural communication. Specifically, EEG hyperconnectivity may indicate a brain signature related to an early stage of cognitive decline possibly leading to dementia [80]. While the EEG-based functional connectivity findings attributable to a specific diagnosis or outcome is far from clear, increased EEG connectivity during the resting state may be a sign of abnormal brain communication, since studies have reported this feature in several neuropsychiatric disorders. For example, individuals with schizophrenia had increased EEG coherence in delta and theta bands relative to controls [81]. Similarly, patients with major depressive disorder exhibited significantly higher EEG coherence as compared to controls in several frequencies, including delta and theta bands [82]. Such alterations in resting-state EEG connectivity in slow rhythms (delta and theta) has also been reported in childhood developmental disorders, such as autism spectrum disorders [83] and specific learning disorders [84]. On the contrary, healthy aging is marked by decreased slow frequency activity (band power) in the delta and theta bands during the resting state [85] as well as by reduced EEG network connectivity [86]. On the other hand, during the task performance, both delta and theta band oscillations predominantly contribute to the generation of P300 or P3 [87], a prominent event-related potential (ERP) component that is a marker of contextual neural processing, the amplitude of which is reduced abnormal in individuals with and/or at risk for AUD, who have shown reduced amplitudes [9]. Interestingly, the slow delta and theta oscillations are often found to be attenuated during task performance in individuals with chronic AUD relative to healthy individuals [88], while these slow theta oscillations are also involved in episodic memory maintenance processes during cognitive processing [89].

At the neural level, it is possible that the hyperconnectivity seen in the *Memory* group may contribute to aberrant synaptic pruning in specific cortical regions [90] in these individuals who have also reported having increased alcohol-related consequences compared to the comparison group. It is also possible that damage to a specific network can enhance connectivity across other regions that are anticorrelated to the damaged network, such as that as it happens in neurodegenerative conditions [91]. Physiologically, alcohol can impact pre- and postsynaptic mechanisms during secretion/recycling of neurotransmitters, leading to the disruption of excitatory and inhibitory neurotransmission [92, 93], potentially caused by detrimental effects of alcohol on glial cells [94]. Recent animal studies confirm that chronic and heavy alcohol consumption can cause aberrant synaptic pruning and substantial loss of excitatory synapses in the prefrontal cortex, resulting in disruption of brain connectivity and dysregulated neural communication across the cortical networks [95]. However, it remains to be confirmed whether the connectivity differences observed in the *Memory* group are the direct consequence of alcohol consumption or indicators of predisposed genetic risk in these individuals, or the interaction of both.

#### 4.1.2. Hyperconnectivity across the hippocampal-cortical networks in the Memory group

Findings reveal that individuals who endorsed alcohol-related memory problems have also shown a predominant pattern of hyperconnectivity across the hippocampal network in their resting EEG, which was recorded about 18 years ago. Specifically, these hyperconnected hippocampal networks (7 out of 8 connections) involved bilateral PHG, bilateral PFC, left LTC, right PCC, and right IPL nodes, spanning delta, theta, and alpha bands [**Fig 7, Panel F**]. Further, majority of the hyper-connected paths (6 out of 7 connections) represented low-frequency (delta/theta) oscillations. Although direct evidence linking EEG-based hyperconnectivity of parahippocampal-cortical network to alcohol-related memory problems is lacking in the literature, some of the available findings may help interpret the findings of the present study. Interestingly, intracranial EEG recordings at the hippocampus and medial temporal regions revealed the existence of independent delta/theta rhythms in different subregions of the human hippocampus and surrounding cortical regions associated with memory encoding and retrieval [96]. Therefore, it is possible that dysregulation (i.e., hyperconnected low frequency paths) in the hippocampal-cortical network, which underlies memory processing [97], may have directly contributed to the alcohol-related memory problems in the *Memory* group. At the neural level, elevated hippocampal resting-state connectivity may be associated with age-related decline in white matter integrity of the fornix as well as deficient neurocognitive function in human adults [98]. Converging findings indicate that memories for recent events underlie dynamic interplay across multiple cortical brain regions and networks, in which the hippocampus acts as a hub integrating information from these subnetworks [99]. Recent studies reveal hippocampal involvement in the default mode network activity. default mode network may mediate interactions between the hippocampus and the neocortex in memory formation and replay [100]. A large neuroimaging study revealed that subregions within default mode network contain fornix fibers from the hippocampus, and thus relating the network to its memory functions [101]. Specifically, the hyperconnected bilateral hippocampal-prefrontal network of slow frequency (delta band) may indicate a dysregulated long-range neural communication involving learning and memory processes, as these networks are crucial for the coordination of activity during memory-guided decision making [102]. Further, the theta band hyperconnectivity of left hippocampal with left temporal cortex and right PCC in *Memory* group may indicate disturbances in verbal [103] and episodic memory [104], respectively. This finding in theta band hippocampal connectivity is important as hippocampal theta rhythm is critical for the optimal functionality of memory networks [105]. It may also be interesting to note that theta band hyperconnectivity across cortical regions was also observed in the APOE-4 carriers of patients with Alzheimer’s disease [106]. Lastly, it needs to be mentioned that a single connection with decreased connectivity at the gamma band in the *Memory* group was observed between ACC and PHG in the right hemisphere. Weaker resting-state connectivity between the hippocampus and ACC may suggest disruption of mood regulation [107], possibly due to compromised structural connectivity between these major structures [108]. Another explanation for lower connectivity between hippocampus and ACC in the *Memory* groups, as it happens in patients with traumatic axonal injury [109], is alcohol-induced microstructural alterations in neuronal fiber tracts connecting brain structures in AUD individuals [110], causing damage to axonal fiber tracts across and within the hemispheres including the hippocampal-cortical bundles [111]. As mentioned earlier, given that the *Memory* group has reported more occasions of heavy drinking and alcohol-related health consequences than the *Control* group, it is expected that neuronal damage, including the compromised hippocampal-cortical connectivity, is more pronounced in these individuals resulting in memory problems along with other neurocognitive and health issues. In sum, it is possible that alcohol-induced hippocampal atrophy [112] may underlie the disruption of cortical hippocampal network subserving memory formation and retrieval processes [113, 114].

#### 4.1.3. Hypoconnectivity across the anterior cingulate hub networks in the Memory group

Findings of the present study have also revealed that the *Memory* group, in addition to the predominant hyperconnectivity across the default mode network nodes in multiple frequencies, manifested six hypoconnected paths (i.e., reduced connectivity strength) across bilateral ACC and other cortical regions (left PFC, bilateral LTC, R.IPL, left PCC, and right PHG) in all frequency bands except the alpha band. All except the connections in the beta band were intra-hemispheric. Broadly, since ACC hub networks within the default mode network are associated with the prediction of outcome for a given choice [115], planning of future actions [116], and social cognition [117], hypoconnectivity of ACC with other cortical regions, including the hippocampal region, may indicate disrupted neural communication leading to less efficient action plans and decision making. ACC also contributes to reward-based action selection or decision-making [118–120] as well as monitoring of action, conflict, error, and outcome [121–124]. In our previous study on EEG source connectivity in abstinent AUD individuals [58], we had also reported hypoconnected prefrontal nodes (PFC and ACC) relaying other cortical regions (LTC, IPL, and PHG) suggesting weaker top-down processing.

Specifically, the hypoconnected ACC–PFC subnetwork in the *Memory* group may suggest compromised top-down cognitive control mediated by the PFC as it happens in individuals addicted to drugs [125]. On the other hand, reduced connectivity of ACC with LTC in the *Memory* group may represent impaired semantic memory processing related to personally relevant action plans in these individuals, as the LTC is related to short-term verbal memory and language processes [126, 127] as well as conceptual representations of actions and behaviors [128, 129]. Further, hypoconnectivity between ACC and IPL in the right hemisphere may indicate a lack of spatial and computational processing for the task at hand, as dictated by the role of right IPL in spatial attention and mathematical cognition [130]. Taken together, these alterations in the brain network may underlie alcohol-induced memory deficits in individuals from the *Memory* group, who have also shown more health problems due to their chronic and/or hazardous alcohol consumption (see Section 4.2. below).

### 4.2. Alcohol Consumption and Health Problems in the Memory group

The top-most predictors of memory problems as revealed by the Random Forests model were alcohol-related consequences during the past 5 years, such as health problems, past negative experiences, withdrawal symptoms, and the largest number of drinks per day. This finding indicates that the individuals with alcohol-related memory problems not only consumed larger quantities of alcohol during the last five years, but also suffered drinking-related adverse consequences such as withdrawal symptoms, negative experiences, and health issues. It is quite possible that the memory problems endorsed by the individuals from the *Memory* group could be one of the health and neurocognitive outcomes due to chronic and/or hazardous alcohol consumption as supported by relevant literature [131–133]. Relatedly, there is also a vast literature documenting alcohol-induced brain damage and cognitive impairments, including memory deficits, in chronic and hazardous drinkers [134–136]. Taken together, alcohol-induced memory problems could be a part of the larger picture of a gross brain damage in chronic and/heavy users of alcohol. Future longitudinal studies combining both structural and functional MRI, along with various EEG and neuropsychological measures, may clarify the exact nature of alcohol-induced neurocognitive deficits.

### 4.3. Personality Features in the Memory group

Among the host of personality and life experience factors included in the Random Forests model, only three factors, namely, harm avoidance, neuroticism, and uplift experiences, were identified as key features that contributed to classifying the *Memory* group from the controls. Our finding suggests increased harm avoidance in the *Memory* group, evidenced by higher endorsement of internalizing traits and negative mood states by these individuals. Although the past studies have shown mixed findings for the harm avoidance subscale of the TPQ in predicting AUD/SUD and risk [38, 137], some of the latter studies have associated these internalizing traits with harmful use of alcohol and other substances [138, 139] and with risk to develop AUD [140–142]. Interestingly, alcohol and other psychoactive substances are often used to self-medicate the negative mood states such as depression [143, 144]. Further, higher neuroticism in the *Memory* group may be related to a variety of alcohol-related outcomes, including relapse [145]. Further, neuroticism has been associated with ineffective use of coping strategies [146], while also mediating the relationship between AUD and neural connectivity [147]. Empirically, neuroticism was has also been found to be associated with internalizing factors related to lifetime diagnosis of mood and anxiety [148]. On the other hand, individuals from the *Memory* group also endorsed fewer uplifting experiences than comparison controls, reflecting less pleasurable experiences at work and home. Lack of adequate uplifting experiences represents a lower buffer against stress and coping [149], which can also contribute to both AUD [146, 150] and internalizing outcomes such as depression [151, 152]. Alternatively, negative mood states may lead to the assessment of fewer experiences as uplifting. Taken together, it is clear that personality and life experience-related factors are important determinants in alcohol-related outcomes, possibly mediated by neural as well as stress-coping dyad mechanisms. However, further studies are necessary to disentangle specific mechanisms involved in the complex etiological pathways of risk, symptoms, and recovery in AUD and related disorders.

### 4.4. Genomic Risk in the Memory group

The only significant PRS measure in the Random Forests model to classifying *Memory* and *Control* groups was derived from the MVP study of DSM-5 AUD, suggesting the importance of AUD-PRS, rather than the consumption related PRS, in predicting neurocognitive outcomes such as alcohol-induced memory problems. This could be partly because individuals from both *Memory* and *Control* groups had a lifetime diagnosis of DSM-IV alcohol dependence. While the DSM-IV alcohol dependence PRS derived from the PGC was not found to be significant, it is possible that it could be because of its relatively smaller GWAS sample size, compared to that of the MVP dataset, and fewer participants of non-European ancestry in the discovery GWAS (see **Table 4**), and/or the more inclusive diagnosis of DSM-5 AUD versus DSM-IV AD.

Nevertheless, the finding that AUD-PRS significantly contributed to the classification suggests that alcohol-induced memory, at least in part, is associated with genomic liability. In general, family studies, twin studies, and GWAS have all demonstrated the heritability of AUD [153–155], and utility of PRS to identify and quantify the risk of developing AUD and related outcomes [65,67,156]. Recently, Lai et al. [67] reported that individuals with AUD had higher PRS than controls and the PRS magnitude increased as the number of DSM-5 diagnostic criteria increased. Further, PRS for alcohol dependence was found to be associated with neural connectivity [36, 157] and cognitive functions, such as verbal fluency, vocabulary, digit-symbol coding, and logical memory [158], as well as brain structure [159]. Unfortunately, PRS related to neurocognitive phenotypes, which could have improved the predictive model, were not included in the study due to a lack of neurocognitive GWAS on AA populations for calculating PRS-CSx for the study sample. Further studies using neurocognitive PRS in multi-ethnic samples are needed to ascertain and quantify the genomic contribution of alcohol-induced memory problems for predictive purposes.

### 4.5. Correlations among the Significant Features

It may be of interest to understand how the significant features, which contributed to the classification of *Memory* individuals from controls, are related to each other. As shown in **Fig. 3**, the correlation matrix revealed some interesting associations. Most obviously, most of the low-frequency connections in the delta and theta frequencies were highly correlated with one another. As mentioned earlier (Section 4.1.2), hippocampal EEG oscillations are mainly represented by delta and theta frequencies, which interact with each other in the memory processes, such as mnemonic encoding and retrieval [96]. Empirically, it is known that delta and theta rhythms are not only correlated with each other but involved in hippocampal-prefrontal communication, which underlies memory and other higher-order cognitive functions such as executive functions [160, 161]. Another interesting finding was that the connections that shared a common node (brain region) between themselves were also significantly correlated with each other, regardless of their frequency band. It is possible that the common node forms a subnetwork that can facilitate information flow across the regions of the subnetwork as well as other connected regions in the brain [162]. Further, correlational results also showed that the beta band connections had highly significant correlations with other connections within the same frequency as well as among low-frequency connections (p < 0.001), especially with the theta band connections (p < 0.001 and survived Bonferroni correction). This could be because low-frequencies (delta/theta) synchronously work together with high-frequencies (beta/gamma) during cognitive processing, including working memory processes [163–165]. However, alpha and gamma band connections showed only within frequency correlations but no cross-frequency correlations, partly because the magnitude of correlations is smaller warranting more statistical power to identify meaningful alpha-gamma associations.

Correlations among the alcohol-related outcome variables were also found to be highly significant with one another, which is in line with the research showing heavy and high-intensity drinking is associated with alcohol-related negative consequences such as withdrawal symptoms and health issues [166, 167]. Further, the significant positive correlation between the two personality traits, namely, neuroticism and harm avoidance, is also backed by the evidence that both traits underlie negative emotions such as fear, shyness, and worry and are regulated by serotonin and opiate pathways [168]. Lastly, it was a rather unexpected finding that there were no highly significant correlations across the domains (e.g., functional connectivity vs. personality), likely because of very low correlation across the domains due to lack of adequate statistical power to detect the subtle associations among features from different categories of predictors.

### 4.6. Limitations and Suggestions

While this is the first multi-modal study including EEG based source connectivity to examine alcohol-related memory problems, which is an important alcohol-related neurocognitive outcome, it has some limitations: (i) the sample size of the study groups is rather small and the findings are therefore only preliminary, (ii) while the groups are matched based on important variables, stratified analyses based on age, sex, and self-reported race, and genetic ancestry, may identify more relevant features specific to each category; (iii) some of the variables were not considered for matching (e.g., memory status during baseline, relatedness among group members, comorbid diagnoses such as substance use, anti-social personality disorder, attention-deficit hyperactivity disorder, etc.), which may have impacted the results; (iv) the memory problems reported by the study sample can be heterogeneous and the assessment of alcohol-related memory problems was only based on oral self-report and not a psychometric measure; studies are currently underway in this sample with comprehensive neurocognitive assessments including memory function and will be more objective and quantitative; (v) the study has not considered genomic or other trait related baseline effects which could have influenced the results, and future large scale studies may consider this aspect into the study design; (vi) recent EEG recordings and neurocognitive assessments, including memory function, in the same sample, which are missing in the current study, but are underway in our lab will further add to predictive modeling; (vii) other specific networks and regions related to memory (e.g., attention and memory networks) have not been explored in the current study, although studies are underway in our lab to explore these networks; (viii) PRS for neurocognitive phenotypes including memory functions have not been included due to lack of availability of multi-ethnic GWAS data. Future studies may attempt to overcome the shortcomings of the study by using a larger sample size and stratified analyses, longitudinal design, multimodal imaging (e.g., fMRI, DTI), and neurocognitive PRS data.

## 5. Conclusions

Our study has elucidated key multimodal features of brain connectivity, personality, life experiences, genomic, and alcohol-related measures that can serve as predictors of later occurring alcohol-related memory problems after about 18 years. Dysregulated brain connectivity, computed from the EEG data collected 18 years ago, in the form of hyper- and hypo-connectivity in specific subnetworks, including the hippocampal-cortical connections, represents potential neural correlates of alcohol-related memory problems. Personality and life experience features such as higher neuroticism and excessive harm avoidance, and fewer uplifting experiences in daily life also contributed to identifying individuals with memory problems from the controls. Importantly, alcohol-related negative consequences during the past 5 years, such as health problems, past negative experiences, withdrawal symptoms, and the largest number of drinks in a day during the past 12 months were the top-most predictors of memory problems. These findings will require confirmation in future studies to: (i) validate these multi-domain features for the use of early identification of individuals who may develop alcohol-induced memory problems in chronic and/or heavy drinkers; and (ii) use EEG-source connectivity measures to further identify/validate specific targets of brain networks underlying AUD related outcomes in general and memory deficits in particular for planning neuromodulation-based treatments (e.g., transcranial magnetic stimulation) as guided by the neural signatures related to dysregulated brain networks in affected individuals. However, in conclusion, the study has many limitations, and the results are only preliminary, warranting large-scale future studies to confirm the current findings by adopting better experimental designs within predictive modeling.

## Acknowledgements

The Collaborative Study on the Genetics of Alcoholism (COGA), Principal Investigators B. Porjesz, V. Hesselbrock, T. Foroud; Scientific Director, A. Agrawal; Translational Director, D. Dick, includes ten different centers: University of Connecticut (V. Hesselbrock); Indiana University (H.J. Edenberg, T. Foroud, Y. Liu, M.H. Plawecki); University of Iowa Carver College of Medicine (S. Kuperman, J. Kramer); SUNY Downstate Health Sciences University (B. Porjesz, J. Meyers, C. Kamarajan, A. Pandey); Washington University in St. Louis (L. Bierut, J. Rice, K. Bucholz, A. Agrawal); University of California at San Diego (M. Schuckit); Rutgers University (J. Tischfield, D. Dick, R. Hart, J. Salvatore); The Children’s Hospital of Philadelphia, University of Pennsylvania (L. Almasy); Icahn School of Medicine at Mount Sinai (A. Goate, P. Slesinger); and Howard University (D. Scott). Other COGA collaborators include: L. Bauer (University of Connecticut); J. Nurnberger Jr., L. Wetherill, X., Xuei, D. Lai, S. O’Connor, (Indiana University); G. Chan (University of Iowa; University of Connecticut); D.B. Chorlian, J. Zhang, P. Barr, S. Kinreich, G. Pandey (SUNY Downstate); N. Mullins (Icahn School of Medicine at Mount Sinai); A. Anokhin, S. Hartz, E. Johnson, V. McCutcheon, S. Saccone (Washington University); J. Moore, F. Aliev, Z. Pang, S. Kuo (Rutgers University); A. Merikangas (The Children’s Hospital of Philadelphia and University of Pennsylvania); H. Chin and A. Parsian are the NIAAA Staff Collaborators. We continue to be inspired by our memories of Henri Begleiter and Theodore Reich, founding PI and Co-PI of COGA, and also owe a debt of gratitude to other past organizers of COGA, including Ting-Kai Li, P. Michael Conneally, Raymond Crowe, and Wendy Reich, for their critical contributions. This national collaborative study is supported by NIH Grant U10AA008401 from the National Institute on Alcohol Abuse and Alcoholism (NIAAA) and the National Institute on Drug Abuse (NIDA).

## Notes

### Competing Interest Statement

The authors have declared no competing interest.

## References

[1] O’Brien CP, McLellan AT (1996) Myths about the treatment of addiction. Lancet, 347(8996), 237–240. https://doi.org/10.1016/s0140-6736(96)90409-2

[2] Koob GF (2014) Neurocircuitry of alcohol addiction: synthesis from animal models. Handb Clin Neurol, 125, 33–54. https://doi.org/10.1016/B978-0-444-62619-6.00003-3

[3] Oscar-Berman M (2000) Neuropsychological vulnerabilities in chronic alcoholism. In: Noronha A, Eckardt MJ, Warren K (eds.): Review of NIAAA’s Neuroscience and Behavioral Research Portfolio. National Institute on Alcohol Abuse and Alcoholism (NIAAA) Research Monograph No. 34, NIAAA, Bethesda, MD, pp. 437–471. https://ia800209.us.archive.org/5/items/reviewofniaaasne00noro/reviewofniaaasne00noro.pdf

[4] Pitel AL, Beaunieux H, Witkowski T, Vabret F, Guillery-Girard B, Quinette P, Desgranges B, Eustache F (2007) Genuine episodic memory deficits and executive dysfunctions in alcoholic subjects early in abstinence. Alcohol Clin Exp Res, 31(7), 1169–1178. https://doi.org/10.1111/j.1530-0277.2007.00418.x

[5] Noel X, Van der Linden M, Brevers D, Campanella S, Hanak C, Kornreich C, Verbanck P (2012) The contribution of executive functions deficits to impaired episodic memory in individuals with alcoholism. Psychiatry Res, 198(1), 116–122. https://doi.org/10.1016/j.psychres.2011.10.007

[6] Oscar-Berman M, Valmas MM, Sawyer KS, Ruiz SM, Luhar RB, Gravitz ZR (2014) Profiles of impaired, spared, and recovered neuropsychologic processes in alcoholism. Handb Clin Neurol, 125, 183–210. https://doi.org/10.1016/B978-0-444-62619-6.00012-4

[7] Le Berre AP, Fama R, Sullivan EV (2017) Executive Functions, Memory, and Social Cognitive Deficits and Recovery in Chronic Alcoholism: A Critical Review to Inform Future Research. Alcohol Clin Exp Res, 41(8), 1432–1443. https://doi.org/10.1111/acer.13431

[8] Enoch MA (2006) Genetic and environmental influences on the development of alcoholism: resilience vs. risk. Ann N Y Acad Sci, 1094, 193–201. https://doi.org/10.1196/annals.1376.019

[9] Porjesz B, Rangaswamy M, Kamarajan C, Jones KA, Padmanabhapillai A, Begleiter H (2005) The utility of neurophysiological markers in the study of alcoholism. Clin Neurophysiol, 116(5), 993–1018. https://doi.org/10.1016/j.clinph.2004.12.016

[10] Miller G, Fagan P (1985) Alcoholism: a polygenic, multifactorial disease. Compr Ther, 11(4), 72–75. https://www.ncbi.nlm.nih.gov/pubmed/4006419

[11] Donner TH, Siegel M (2011) A framework for local cortical oscillation patterns. Trends Cogn Sci, 15(5), 191–199. https://doi.org/10.1016/j.tics.2011.03.007

[12] Duzel E, Penny WD, Burgess N (2010) Brain oscillations and memory. Curr Opin Neurobiol, 20(2), 143–149. https://doi.org/10.1016/j.conb.2010.01.004

[13] Buzsaki G, Moser EI (2013) Memory, navigation and theta rhythm in the hippocampal-entorhinal system. Nat Neurosci, 16(2), 130–138. https://doi.org/10.1038/nn.3304

[14] Klimesch W (1996) Memory processes, brain oscillations and EEG synchronization. Int J Psychophysiol, 24(1-2), 61–100. https://doi.org/10.1016/s0167-8760(96)00057-8

[15] Hanslmayr S, Staudigl T (2014) How brain oscillations form memories--a processing based perspective on oscillatory subsequent memory effects. Neuroimage, 85 Pt 2, 648–655. https://doi.org/10.1016/j.neuroimage.2013.05.121

[16] Kahana MJ (2006) The cognitive correlates of human brain oscillations. J Neurosci, 26(6), 1669–1672. https://doi.org/10.1523/JNEUROSCI.3737-05c.2006

[17] Herweg NA, Solomon EA, Kahana MJ (2020) Theta Oscillations in Human Memory. Trends Cogn Sci, 24(3), 208–227. https://doi.org/10.1016/j.tics.2019.12.006

[18] Colgin LL (2011) Oscillations and hippocampal-prefrontal synchrony. Curr Opin Neurobiol, 21(3), 467–474. https://doi.org/10.1016/j.conb.2011.04.006

[19] Colgin LL (2013) Mechanisms and functions of theta rhythms. Annu Rev Neurosci, 36, 295–312. https://doi.org/10.1146/annurev-neuro-062012-170330

[20] Colgin LL (2015) Theta-gamma coupling in the entorhinal-hippocampal system. Curr Opin Neurobiol, 31, 45–50. https://doi.org/10.1016/j.conb.2014.08.001

[21] Canolty RT, Knight RT (2010) The functional role of cross-frequency coupling. Trends Cogn Sci, 14(11), 506–515. https://doi.org/10.1016/j.tics.2010.09.001

[22] Pascual-Marqui RD, Lehmann D, Koukkou M, Kochi K, Anderer P, Saletu B, Tanaka H, Hirata K, John ER, Prichep L, Biscay-Lirio R, Kinoshita T (2011) Assessing interactions in the brain with exact low-resolution electromagnetic tomography. Philos Trans A Math Phys Eng Sci, 369(1952), 3768–3784. https://doi.org/10.1098/rsta.2011.0081

[23] Canuet L, Ishii R, Pascual-Marqui RD, Iwase M, Kurimoto R, Aoki Y, Ikeda S, Takahashi H, Nakahachi T, Takeda M (2011) Resting-state EEG source localization and functional connectivity in schizophrenia-like psychosis of epilepsy. PLOS ONE, 6(11), e27863. https://doi.org/10.1371/journal.pone.0027863

[24] Bowyer SM (2016) Coherence a measure of the brain networks: past and present. Neuropsychiatric Electrophysiology, 2(1), 1–12. https://doi.org/10.1186/s40810-015-0015-7

[25] Nolte G, Bai O, Wheaton L, Mari Z, Vorbach S, Hallett M (2004) Identifying true brain interaction from EEG data using the imaginary part of coherency. Clin Neurophysiol, 115(10), 2292–2307. https://doi.org/10.1016/j.clinph.2004.04.029

[26] Olbrich S, Trankner A, Chittka T, Hegerl U, Schonknecht P (2014) Functional connectivity in major depression: Increased phase synchronization between frontal cortical EEG-source estimates. Psychiatry Res, 222(1-2), 91–99. https://doi.org/10.1016/j.pscychresns.2014.02.010

[27] Hata M, Kazui H, Tanaka T, Ishii R, Canuet L, Pascual-Marqui RD, Aoki Y, Ikeda S, Kanemoto H, Yoshiyama K, Iwase M, Takeda M (2016) Functional connectivity assessed by resting state EEG correlates with cognitive decline of Alzheimer’s disease - An eLORETA study. Clin Neurophysiol, 127(2), 1269–1278. https://doi.org/10.1016/j.clinph.2015.10.030

[28] Imperatori C, Della Marca G, Brunetti R, Carbone GA, Massullo C, Valenti EM, Amoroso N, Maestoso G, Contardi A, Farina B (2016) Default Mode Network alterations in alexithymia: an EEG power spectra and connectivity study. Sci Rep, 6, 36653. https://doi.org/10.1038/srep36653

[29] Whitton AE, Deccy S, Ironside ML, Kumar P, Beltzer M, Pizzagalli DA (2018) Electroencephalography Source Functional Connectivity Reveals Abnormal High-Frequency Communication Among Large-Scale Functional Networks in Depression. Biol Psychiatry Cogn Neurosci Neuroimaging, 3(1), 50–58. https://doi.org/10.1016/j.bpsc.2017.07.001

[30] Huang Y, Mohan A, De Ridder D, Sunaert S, Vanneste S (2018) The neural correlates of the unified percept of alcohol-related craving: a fMRI and EEG study. Sci Rep, 8(1), 923. https://doi.org/10.1038/s41598-017-18471-y

[31] Huijbers W, Pennartz CM, Cabeza R, Daselaar SM (2011) The hippocampus is coupled with the default network during memory retrieval but not during memory encoding. PLOS ONE, 6(4), e17463. https://doi.org/10.1371/journal.pone.0017463

[32] Westlye ET, Lundervold A, Rootwelt H, Lundervold AJ, Westlye LT (2011) Increased hippocampal default mode synchronization during rest in middle-aged and elderly APOE epsilon4 carriers: relationships with memory performance. J Neurosci, 31(21), 7775–7783. https://doi.org/10.1523/JNEUROSCI.1230-11.2011

[33] Ward AM, Schultz AP, Huijbers W, Van Dijk KR, Hedden T, Sperling RA (2013) The parahippocampal gyrus links the default-mode cortical network with the medial temporal lobe memory system. Hum Brain Mapp. https://doi.org/10.1002/hbm.22234

[34] Huo L, Li R, Wang P, Zheng Z, Li J (2018) The Default Mode Network Supports Episodic Memory in Cognitively Unimpaired Elderly Individuals: Different Contributions to Immediate Recall and Delayed Recall. Front Aging Neurosci, 10(6), 6. https://doi.org/10.3389/fnagi.2018.00006

[35] Nurnberger JI, Jr., Wang Y, Zang Y, Lai D, Wetherill L, Edenberg HJ, Aliev F, Plawecki MH, Chorlian D, Chan G, Bucholz K, Bauer L, Kamarajan C, Salvatore JE, Kapoor M, Hesselbrock V, Dick D, Bierut L, McCutcheon V, Meyers JL, Porjesz B, Kramer J, Kuperman S, Kinreich S, Anokhin AP, Collaborative Study on the Genetics of A (2022) High Polygenic Risk Scores Are Associated With Early Age of Onset of Alcohol Use Disorder in Adolescents and Young Adults at Risk. Biol Psychiatry Glob Open Sci, 2(4), 379–388. https://doi.org/10.1016/j.bpsgos.2021.10.007

[36] Kinreich S, McCutcheon VV, Aliev F, Meyers JL, Kamarajan C, Pandey AK, Chorlian DB, Zhang J, Kuang W, Pandey G, Viteri SS, Francis MW, Chan G, Bourdon JL, Dick DM, Anokhin AP, Bauer L, Hesselbrock V, Schuckit MA, Nurnberger JI, Jr., Foroud TM, Salvatore JE, Bucholz KK, Porjesz B (2021) Predicting alcohol use disorder remission: a longitudinal multimodal multi-featured machine learning approach. Transl Psychiatry, 11(1), 166. https://doi.org/10.1038/s41398-021-01281-2

[37] Li JJ, Savage JE, Kendler KS, Hickman M, Mahedy L, Macleod J, Kaprio J, Rose RJ, Dick DM (2017) Polygenic Risk, Personality Dimensions, and Adolescent Alcohol Use Problems: A Longitudinal Study. J Stud Alcohol Drugs, 78(3), 442–451. https://doi.org/10.15288/jsad.2017.78.442

[38] Yoshino A, Kato M, Takeuchi M, Ono Y, Kitamura T (1994) Examination of the tridimensional personality hypothesis of alcoholism using empirically multivariate typology. Alcohol Clin Exp Res, 18(5), 1121–1124. https://doi.org/10.1111/j.1530-0277.1994.tb00091.x

[39] Tomassini A, Struglia F, Spaziani D, Pacifico R, Stratta P, Rossi A (2012) Decision making, impulsivity, and personality traits in alcohol-dependent subjects. Am J Addict, 21(3), 263–267. https://doi.org/10.1111/j.1521-0391.2012.00225.x

[40] Littlefield AK, Sher KJ (2010) The Multiple, Distinct Ways that Personality Contributes to Alcohol Use Disorders. Soc Personal Psychol Compass, 4(9), 767–782. https://doi.org/10.1111/j.1751-9004.2010.00296.x

[41] Rosenstrom T, Torvik FA, Ystrom E, Czajkowski NO, Gillespie NA, Aggen SH, Krueger RF, Kendler KS, Reichborn-Kjennerud T (2018) Prediction of alcohol use disorder using personality disorder traits: a twin study. Addiction, 113(1), 15–24. https://doi.org/10.1111/add.13951

[42] Creed M (2018) Current and emerging neuromodulation therapies for addiction: insight from pre-clinical studies. Curr Opin Neurobiol, 49, 168–174. https://doi.org/10.1016/j.conb.2018.02.015

[43] Reinhart RMG, Nguyen JA (2019) Working memory revived in older adults by synchronizing rhythmic brain circuits. Nat Neurosci, 22(5), 820–827. https://doi.org/10.1038/s41593-019-0371-x

[44] Schuckit MA, Smith TL, Danko G, Kramer J, Bucholz KK, McCutcheon V, Chan G, Kuperman S, Hesselbrock V, Dick DM, Hesselbrock M, Porjesz B, Edenberg HJ, Nureberger JI, Jr., Gregg M, Schoen L, Kawamura M, Mendoza LA (2018) A 22-Year Follow-Up (Range 16 to 23) of Original Subjects with Baseline Alcohol Use Disorders from the Collaborative Study on Genetics of Alcoholism. Alcohol Clin Exp Res, 42(9), 1704–1714. https://doi.org/10.1111/acer.13810

[45] Chan G, Kramer JR, Schuckit MA, Hesselbrock V, Bucholz KK, Edenberg HJ, Acion L, Langbehn D, McCutcheon V, Nurnberger JI, Jr., Hesselbrock M, Porjesz B, Bierut L, Marenna BC, Cookman A, Kuperman S (2019) A Pilot Follow-Up Study of Older Alcohol-Dependent COGA Adults. Alcohol Clin Exp Res, 43(8), 1759–1768. https://doi.org/10.1111/acer.14116

[46] Begleiter H, Reich T, Hesselbrock V, Porjesz B, Li T-K, Schuckit MA, Edenberg HJ, Rice JP (1995) The Collaborative Study on the Genetics of Alcoholism. Alcohol Health Res World, 19(2), 228–236. https://pubmed.ncbi.nlm.nih.gov/31798102/

[47] Bucholz KK, McCutcheon VV, Agrawal A, Dick DM, Hesselbrock VM, Kramer JR, Kuperman S, Nurnberger JI, Jr., Salvatore JE, Schuckit MA, Bierut LJ, Foroud TM, Chan G, Hesselbrock M, Meyers JL, Edenberg HJ, Porjesz B (2017) Comparison of Parent, Peer, Psychiatric, and Cannabis Use Influences Across Stages of Offspring Alcohol Involvement: Evidence from the COGA Prospective Study. Alcohol Clin Exp Res, 41(2), 359–368. https://doi.org/10.1111/acer.13293

[48] Lai D, Wetherill L, Bertelsen S, Carey CE, Kamarajan C, Kapoor M, Meyers JL, Anokhin AP, Bennett DA, Bucholz KK, Chang KK, De Jager PL, Dick DM, Hesselbrock V, Kramer J, Kuperman S, Nurnberger JI, Jr., Raj T, Schuckit M, Scott DM, Taylor RE, Tischfield J, Hariri AR, Edenberg HJ, Agrawal A, Bogdan R, Porjesz B, Goate AM, Foroud T (2019) Genome-wide association studies of alcohol dependence, DSM-IV criterion count and individual criteria. Genes Brain Behav, 18(6), e12579. https://doi.org/10.1111/gbb.12579

[49] Bucholz KK, Cadoret R, Cloninger CR, Dinwiddie SH, Hesselbrock VM, Nurnberger JI, Jr., Reich T, Schmidt I, Schuckit MA (1994) A new, semi-structured psychiatric interview for use in genetic linkage studies: a report on the reliability of the SSAGA. J Stud Alcohol, 55(2), 149–158. https://doi.org/10.15288/jsa.1994.55.149

[50] Hesselbrock M, Easton C, Bucholz KK, Schuckit M, Hesselbrock V (1999) A validity study of the SSAGA--a comparison with the SCAN. Addiction, 94(9), 1361–1370. https://doi.org/10.1046/j.1360-0443.1999.94913618.x

[51] Harris PA, Taylor R, Minor BL, Elliott V, Fernandez M, O’Neal L, McLeod L, Delacqua G, Delacqua F, Kirby J, Duda SN, REDCap Consortium (2019) The REDCap consortium: Building an international community of software platform partners. J Biomed Inform, 95, 103208. https://doi.org/10.1016/j.jbi.2019.103208

[52] Harris PA, Taylor R, Thielke R, Payne J, Gonzalez N, Conde JG (2009) Research electronic data capture (REDCap)--a metadata-driven methodology and workflow process for providing translational research informatics support. J Biomed Inform, 42(2), 377–381. https://doi.org/10.1016/j.jbi.2008.08.010

[53] Rangaswamy M, Porjesz B, Chorlian DB, Wang K, Jones KA, Bauer LO, Rohrbaugh J, O’Connor SJ, Kuperman S, Reich T, Begleiter H (2002) Beta power in the EEG of alcoholics. Biol Psychiatry, 52(8), 831–842. https://doi.org/10.1016/s0006-3223(02)01362-8

[54] Chorlian DB, Rangaswamy M, Porjesz B (2009) EEG coherence: topography and frequency structure. Exp Brain Res, 198(1), 59–83. https://doi.org/10.1007/s00221-009-1936-9

[55] Pascual-Marqui RD (2007) Discrete, 3D distributed, linear imaging methods of electric neuronal activity. Part 1: exact, zero error localization. *arXiv*, 0710.3341. http://arxiv.org/pdf/0710.3341

[56] Buckner RL, Andrews-Hanna JR, Schacter DL (2008) The brain’s default network: anatomy, function, and relevance to disease. Ann N Y Acad Sci, 1124, 1–38. https://doi.org/10.1196/annals.1440.011

[57] Thatcher RW, North DM, Biver CJ (2014) LORETA EEG phase reset of the default mode network. Front Hum Neurosci, 8, 529. https://doi.org/10.3389/fnhum.2014.00529

[58] Kamarajan C, Ardekani BA, Pandey AK, Chorlian DB, Kinreich S, Pandey G, Meyers JL, Zhang J, Kuang W, Stimus AT, Porjesz B (2020) Random Forest Classification of Alcohol Use Disorder Using EEG Source Functional Connectivity, Neuropsychological Functioning, and Impulsivity Measures. Behav Sci (Basel), 10(3), 62. https://doi.org/10.3390/bs10030062

[59] Kamarajan C, Ardekani BA, Pandey AK, Kinreich S, Pandey G, Chorlian DB, Meyers JL, Zhang J, Bermudez E, Stimus AT, Porjesz B (2020) Random Forest Classification of Alcohol Use Disorder Using fMRI Functional Connectivity, Neuropsychological Functioning, and Impulsivity Measures. Brain Sci, 10(2), 115. https://doi.org/10.3390/brainsci10020115

[60] Kranzler HR, Zhou H, Kember RL, Vickers Smith R, Justice AC, Damrauer S, Tsao PS, Klarin D, Baras A, Reid J, Overton J, Rader DJ, Cheng Z, Tate JP, Becker WC, Concato J, Xu K, Polimanti R, Zhao H, Gelernter J (2019) Genome-wide association study of alcohol consumption and use disorder in 274,424 individuals from multiple populations. Nat Commun, 10(1), 1499. https://doi.org/10.1038/s41467-019-09480-8

[61] Gelernter J, Sun N, Polimanti R, Pietrzak RH, Levey DF, Lu Q, Hu Y, Li B, Radhakrishnan K, Aslan M, Cheung KH, Li Y, Rajeevan N, Sayward F, Harrington K, Chen Q, Cho K, Honerlaw J, Pyarajan S, Lencz T, Quaden R, Shi Y, Hunter-Zinck H, Gaziano JM, Kranzler HR, Concato J, Zhao H, Stein MB, Department of Veterans Affairs Cooperative Studies P, Million Veteran P (2019) Genome-wide Association Study of Maximum Habitual Alcohol Intake in >140,000 U.S. European and African American Veterans Yields Novel Risk Loci. Biol Psychiatry, 86(5), 365-376. https://doi.org/10.1016/j.biopsych.2019.03.984

[62] Walters RK, Polimanti R, Johnson EC, McClintick JN, Adams MJ, Adkins AE, Aliev F, Bacanu SA, Batzler A, Bertelsen S, Biernacka JM, Bigdeli TB, Chen LS, Clarke TK, Chou YL, Degenhardt F, Docherty AR, Edwards AC, Fontanillas P, Foo JC, Fox L, Frank J, Giegling I, Gordon S, Hack LM, Hartmann AM, Hartz SM, Heilmann-Heimbach S, Herms S, Hodgkinson C, Hoffmann P, Jan Hottenga J, Kennedy MA, Alanne-Kinnunen M, Konte B, Lahti J, Lahti-Pulkkinen M, Lai D, Ligthart L, Loukola A, Maher BS, Mbarek H, McIntosh AM, McQueen MB, Meyers JL, Milaneschi Y, Palviainen T, Pearson JF, Peterson RE, Ripatti S, Ryu E, Saccone NL, Salvatore JE, Sanchez-Roige S, Schwandt M, Sherva R, Streit F, Strohmaier J, Thomas N, Wang JC, Webb BT, Wedow R, Wetherill L, Wills AG, and Me Research T, Boardman JD, Chen D, Choi DS, Copeland WE, Culverhouse RC, Dahmen N, Degenhardt L, Domingue BW, Elson SL, Frye MA, Gabel W, Hayward C, Ising M, Keyes M, Kiefer F, Kramer J, Kuperman S, Lucae S, Lynskey MT, Maier W, Mann K, Mannisto S, Muller-Myhsok B, Murray AD, Nurnberger JI, Palotie A, Preuss U, Raikkonen K, Reynolds MD, Ridinger M, Scherbaum N, Schuckit MA, Soyka M, Treutlein J, Witt S, Wodarz N, Zill P, Adkins DE, Boden JM, Boomsma DI, Bierut LJ, Brown SA, Bucholz KK, Cichon S, Costello EJ, de Wit H, Diazgranados N, Dick DM, Eriksson JG, Farrer LA, Foroud TM, Gillespie NA, Goate AM, Goldman D, Grucza RA, Hancock DB, Harris KM, Heath AC, Hesselbrock V, Hewitt JK, Hopfer CJ, Horwood J, Iacono W, Johnson EO, Kaprio JA, Karpyak VM, Kendler KS, Kranzler HR, Krauter K, Lichtenstein P, Lind PA, McGue M, MacKillop J, Madden PAF, Maes HH, Magnusson P, Martin NG, Medland SE, Montgomery GW, Nelson EC, Nothen MM, Palmer AA, Pedersen NL, Penninx B, Porjesz B, Rice JP, Rietschel M, Riley BP, Rose R, Rujescu D, Shen PH, Silberg J, Stallings MC, Tarter RE, Vanyukov MM, Vrieze S, Wall TL, Whitfield JB, Zhao H, Neale BM, Gelernter J, Edenberg HJ, Agrawal A (2018) Transancestral GWAS of alcohol dependence reveals common genetic underpinnings with psychiatric disorders. Nat Neurosci, 21(12), 1656–1669. https://doi.org/10.1038/s41593-018-0275-1

[63] Ge T, Chen CY, Ni Y, Feng YA, Smoller JW (2019) Polygenic prediction via Bayesian regression and continuous shrinkage priors. Nat Commun, 10(1), 1776. https://doi.org/10.1038/s41467-019-09718-5

[64] Ruan Y, Lin Y-F, Feng Y-CA, Chen C-Y, Lam M, Guo Z, He L, Sawa A, Martin AR, Qin S, Huang H, Ge T (2021) Improving Polygenic Prediction in Ancestrally Diverse Populations. medRxiv, 2020.2012.2027.20248738. https://doi.org/10.1101/2020.12.27.20248738

[65] Barr PB, Ksinan A, Su J, Johnson EC, Meyers JL, Wetherill L, Latvala A, Aliev F, Chan G, Kuperman S, Nurnberger J, Kamarajan C, Anokhin A, Agrawal A, Rose RJ, Edenberg HJ, Schuckit M, Kaprio J, Dick DM (2020) Using polygenic scores for identifying individuals at increased risk of substance use disorders in clinical and population samples. Transl Psychiatry, 10(1), 196. https://doi.org/10.1038/s41398-020-00865-8

[66] Lai D, Schwantes-An TH, Abreu M, Chan G, Hesselbrock V, Kamarajan C, Liu Y, Meyers JL, Nurnberger JI, Jr., Plawecki MH, Wetherill L, Schuckit M, Zhang P, Edenberg HJ, Porjesz B, Agrawal A, Foroud T (2022) Gene-based polygenic risk scores analysis of alcohol use disorder in African Americans. Transl Psychiatry, 12(1), 266. https://doi.org/10.1038/s41398-022-02029-2

[67] Lai D, Johnson EC, Colbert S, Pandey G, Chan G, Bauer L, Francis MW, Hesselbrock V, Kamarajan C, Kramer J, Kuang W, Kuo S, Kuperman S, Liu Y, McCutcheon V, Pang Z, Plawecki MH, Schuckit M, Tischfield J, Wetherill L, Zang Y, Edenberg HJ, Porjesz B, Agrawal A, Foroud T (2022) Evaluating risk for alcohol use disorder: Polygenic risk scores and family history. Alcohol Clin Exp Res, 46(3), 374–383. https://doi.org/10.1111/acer.14772

[68] Ge T, Irvin MR, Patki A, Srinivasasainagendra V, Lin YF, Tiwari HK, Armstrong ND, Benoit B, Chen CY, Choi KW, Cimino JJ, Davis BH, Dikilitas O, Etheridge B, Feng YA, Gainer V, Huang H, Jarvik GP, Kachulis C, Kenny EE, Khan A, Kiryluk K, Kottyan L, Kullo IJ, Lange C, Lennon N, Leong A, Malolepsza E, Miles AD, Murphy S, Namjou B, Narayan R, O’Connor MJ, Pacheco JA, Perez E, Rasmussen-Torvik LJ, Rosenthal EA, Schaid D, Stamou M, Udler MS, Wei WQ, Weiss ST, Ng MCY, Smoller JW, Lebo MS, Meigs JB, Limdi NA, Karlson EW (2022) Development and validation of a trans-ancestry polygenic risk score for type 2 diabetes in diverse populations. Genome Med, 14(1), 70. https://doi.org/10.1186/s13073-022-01074-2

[69] Kamarajan C, Ardekani BA, Pandey AK, Kinreich S, Pandey G, Chorlian DB, Meyers JL, Zhang J, Bermudez E, Kuang W, Stimus AT, Porjesz B (2022) Differentiating Individuals with and without Alcohol Use Disorder Using Resting-State fMRI Functional Connectivity of Reward Network, Neuropsychological Performance, and Impulsivity Measures. Behav Sci (Basel), 12(5), 128. https://doi.org/10.3390/bs12050128

[70] Nguyen C, Wang Y, Nguyen HN (2013) Random forest classifier combined with feature selection for breast cancer diagnosis and prognostic. J Biomed Sci Eng, 06(05), 551–560. https://doi.org/10.4236/jbise.2013.65070

[71] Chandrashekar G, Sahin F (2014) A survey on feature selection methods. Computers & Electrical Engineering, 40(1), 16–28. https://doi.org/10.1016/j.compeleceng.2013.11.024

[72] Cai J, Luo JW, Wang SL, Yang S (2018) Feature selection in machine learning: A new perspective. Neurocomputing, 300, 70–79. https://doi.org/10.1016/j.neucom.2017.11.077

[73] Kamala RF, Thangaiah PRJ (2019) A Novel Two-Stage Selection of Feature Subsets in Machine Learning. Eng Technol Appl Sci Res, 9(3), 4169–4175. https://doi.org/10.48084/etasr.2735

[74] Raj S, Singh S, Kumar A, Sarkar S, Pradhan C (2021) Feature Selection and Random Forest Classification for Breast Cancer Disease. In: Data Analytics in Bioinformatics, Wiley, Hoboken, NJ, USA, pp. 191–210. https://doi.org/10.1002/9781119785620.ch8

[75] Liaw A, Wiener M (2018) Breiman and Cutler’s Random Forests for Classification and Regression. https://cran.r-project.org/web/packages/randomForest/

[76] Kuhn M, Wing J, Weston S, Williams A, Keefer C, Engelhardt A, Cooper T, Mayer Z, Kenkel B, Benesty M, Lescarbeau R, Ziem A, Scrucca L, Tang Y, Candan C, Hunt T (2019) *Classification and Regression Training.* R Package Version 6.0-84, URL: https://cran.r-project.org/web/packages/caret

[77] Paluszynska A, Biecek P, Jiang Y (2019) randomForestExplainer: Explaining and Visualizing Random Forests in Terms of Variable Importance. R Package Version 0.10.0, URL: https://cran.r-project.org/web/packages/randomForestExplainer

[78] Semon R (1921) The Mneme. G. Allen & Unwin Limited, London. https://books.google.com/books?hl=en&lr=&id=I0uQ6yzHxPIC&oi=fnd&pg=PA9&dq=Semon++(1921)+The+Mneme&ots=WztcQkPJqR&sig=GGUK6P7uLjrnwh1vrbmFbwJ0iIk#v=onepage&q=Semon%20R%20(1921)%20The%20Mneme&f=false

[79] Hebb DO (1949) The organization of behavior: A neuropsychological theory. John Wiley & Sons, Inc, New York. https://doi.org/10.4324/9781410612403

[80] Bonanni L, Moretti D, Benussi A, Ferri L, Russo M, Carrarini C, Barbone F, Arnaldi D, Falasca NW, Koch G, Cagnin A, Nobili F, Babiloni C, Borroni B, Padovani A, Onofrj M, Franciotti R, group-SINDEM FTDIs (2021) Hyperconnectivity in Dementia Is Early and Focal and Wanes with Progression. Cereb Cortex, 31(1), 97–105. https://doi.org/10.1093/cercor/bhaa209

[81] Di Lorenzo G, Daverio A, Ferrentino F, Santarnecchi E, Ciabattini F, Monaco L, Lisi G, Barone Y, Di Lorenzo C, Niolu C, Seri S, Siracusano A (2015) Altered resting-state EEG source functional connectivity in schizophrenia: the effect of illness duration. Front Hum Neurosci, 9, 234. https://doi.org/10.3389/fnhum.2015.00234

[82] Leuchter AF, Cook IA, Hunter AM, Cai C, Horvath S (2012) Resting-state quantitative electroencephalography reveals increased neurophysiologic connectivity in depression. PLOS ONE, 7(2), e32508. https://doi.org/10.1371/journal.pone.0032508

[83] Wang J, Wang X, Wang X, Zhang H, Zhou Y, Chen L, Li Y, Wu L (2020) Increased EEG coherence in long-distance and short-distance connectivity in children with autism spectrum disorders. Brain Behav, 10(10), e01796. https://doi.org/10.1002/brb3.1796

[84] Arns M, Peters S, Breteler R, Verhoeven L (2007) Different brain activation patterns in dyslexic children: evidence from EEG power and coherence patterns for the double-deficit theory of dyslexia. J Integr Neurosci, 6(1), 175–190. https://doi.org/10.1142/s0219635207001404

[85] Vlahou EL, Thurm F, Kolassa IT, Schlee W (2014) Resting-state slow wave power, healthy aging and cognitive performance. Sci Rep, 4(1), 5101. https://doi.org/10.1038/srep05101

[86] Javaid H, Kumarnsit E, Chatpun S (2022) Age-Related Alterations in EEG Network Connectivity in Healthy Aging. Brain Sci, 12(2). https://doi.org/10.3390/brainsci12020218

[87] Basar-Eroglu C, Basar E, Demiralp T, Schurmann M (1992) P300-response: possible psychophysiological correlates in delta and theta frequency channels. A review. Int J Psychophysiol, 13(2), 161–179. http://www.ncbi.nlm.nih.gov/pubmed/1399755

[88] Pandey AK, Kamarajan C, Rangaswamy M, Porjesz B (2012) Event-Related Oscillations in Alcoholism Research: A Review. *J Addict Res Ther*, Suppl 7(1), 1–13. https://doi.org/10.4172/2155-6105.S7-001

[89] Toth B, Boha R, Posfai M, Gaal ZA, Konya A, Stam CJ, Molnar M (2012) EEG synchronization characteristics of functional connectivity and complex network properties of memory maintenance in the delta and theta frequency bands. Int J Psychophysiol, 83(3), 399–402. https://doi.org/10.1016/j.ijpsycho.2011.11.017

[90] Sakai J (2020) Core Concept: How synaptic pruning shapes neural wiring during development and, possibly, in disease. Proc Natl Acad Sci U S A, 117(28), 16096–16099. https://doi.org/10.1073/pnas.2010281117

[91] Pievani M, de Haan W, Wu T, Seeley WW, Frisoni GB (2011) Functional network disruption in the degenerative dementias. Lancet Neurol, 10(9), 829–843. https://doi.org/10.1016/S1474-4422(11)70158-2

[92] Lovinger DM, Roberto M (2013) Synaptic effects induced by alcohol. Curr Top Behav Neurosci, 13, 31–86. https://doi.org/10.1007/7854_2011_143

[93] Abrahao KP, Salinas AG, Lovinger DM (2017) Alcohol and the Brain: Neuronal Molecular Targets, Synapses, and Circuits. Neuron, 96(6), 1223–1238. https://doi.org/10.1016/j.neuron.2017.10.032

[94] Lacagnina MJ, Rivera PD, Bilbo SD (2017) Glial and Neuroimmune Mechanisms as Critical Modulators of Drug Use and Abuse. Neuropsychopharmacology, 42(1), 156–177. https://doi.org/10.1038/npp.2016.121

[95] Socodato R, Henriques JF, Portugal CC, Almeida TO, Tedim-Moreira J, Alves RL, Canedo T, Silva C, Magalhaes A, Summavielle T, Relvas JB (2020) Daily alcohol intake triggers aberrant synaptic pruning leading to synapse loss and anxiety-like behavior. Sci Signal, 13(650). https://doi.org/10.1126/scisignal.aba5754

[96] Mormann F, Osterhage H, Andrzejak RG, Weber B, Fernandez G, Fell J, Elger CE, Lehnertz K (2008) Independent delta/theta rhythms in the human hippocampus and entorhinal cortex. Front Hum Neurosci, 2, 3. https://doi.org/10.3389/neuro.09.003.2008

[97] Inhoff MC, Ranganath C (2017) Dynamic Cortico-hippocampal Networks Underlying Memory and Cognition: The PMAT Framework. In: Hannula DE, Duff MC (eds.): The Hippocampus from Cells to Systems, Springer International Publishing, Cham, pp. 559–589. https://doi.org/10.1007/978-3-319-50406-3_18

[98] Salami A, Pudas S, Nyberg L (2014) Elevated hippocampal resting-state connectivity underlies deficient neurocognitive function in aging. Proc Natl Acad Sci U S A, 111(49), 17654–17659. https://doi.org/10.1073/pnas.1410233111

[99] Schedlbauer AM, Copara MS, Watrous AJ, Ekstrom AD (2014) Multiple interacting brain areas underlie successful spatiotemporal memory retrieval in humans. Sci Rep, 4(1), 6431. https://doi.org/10.1038/srep06431

[100] Kaefer K, Stella F, McNaughton BL, Battaglia FP (2022) Replay, the default mode network and the cascaded memory systems model. Nat Rev Neurosci, 23(10), 628–640. https://doi.org/10.1038/s41583-022-00620-6

[101] Kernbach JM, Yeo BTT, Smallwood J, Margulies DS, Thiebaut de Schotten M, Walter H, Sabuncu MR, Holmes AJ, Gramfort A, Varoquaux G, Thirion B, Bzdok D (2018) Subspecialization within default mode nodes characterized in 10,000 UK Biobank participants. Proc Natl Acad Sci U S A, 115(48), 12295–12300. https://doi.org/10.1073/pnas.1804876115

[102] Shin JD, Jadhav SP (2016) Multiple modes of hippocampal-prefrontal interactions in memory-guided behavior. Curr Opin Neurobiol, 40, 161–169. https://doi.org/10.1016/j.conb.2016.07.015

[103] Kucewicz MT, Berry BM, Miller LR, Khadjevand F, Ezzyat Y, Stein JM, Kremen V, Brinkmann BH, Wanda P, Sperling MR, Gorniak R, Davis KA, Jobst BC, Gross RE, Lega B, Van Gompel J, Stead SM, Rizzuto DS, Kahana MJ, Worrell GA (2018) Evidence for verbal memory enhancement with electrical brain stimulation in the lateral temporal cortex. Brain, 141(4), 971–978. https://doi.org/10.1093/brain/awx373

[104] McCormick C, Protzner AB, Barnett AJ, Cohn M, Valiante TA, McAndrews MP (2014) Linking DMN connectivity to episodic memory capacity: what can we learn from patients with medial temporal lobe damage? Neuroimage Clin, 5, 188–196. https://doi.org/10.1016/j.nicl.2014.05.008

[105] Nicolas B, Sala-Padro J, Cucurell D, Santurino M, Falip M, Fuentemilla L (2021) Theta rhythm supports hippocampus-dependent integrative encoding in schematic/semantic memory networks. Neuroimage, 226, 117558. https://doi.org/10.1016/j.neuroimage.2020.117558

[106] Canuet L, Tellado I, Couceiro V, Fraile C, Fernandez-Novoa L, Ishii R, Takeda M, Cacabelos R (2012) Resting-state network disruption and APOE genotype in Alzheimer’s disease: a lagged functional connectivity study. PLOS ONE, 7(9), e46289. https://doi.org/10.1371/journal.pone.0046289

[107] Zhang W, Liu X, Zhang Y, Song L, Hou J, Chen B, He M, Cai P, Lii H (2014) Disrupted functional connectivity of the hippocampus in patients with hyperthyroidism: evidence from resting-state fMRI. Eur J Radiol, 83(10), 1907–1913. https://doi.org/10.1016/j.ejrad.2014.07.003

[108] Fani N, King TZ, Shin J, Srivastava A, Brewster RC, Jovanovic T, Bradley B, Ressler KJ (2016) Structural and Functional Connectivity in Posttraumatic Stress Disorder: Associations with Fkbp5. Depress Anxiety, 33(4), 300–307. https://doi.org/10.1002/da.22483

[109] Marquez de la Plata CD, Garces J, Shokri Kojori E, Grinnan J, Krishnan K, Pidikiti R, Spence J, Devous MD, Sr., Moore C, McColl R, Madden C, Diaz-Arrastia R (2011) Deficits in functional connectivity of hippocampal and frontal lobe circuits after traumatic axonal injury. Arch Neurol, 68(1), 74–84. https://doi.org/10.1001/archneurol.2010.342

[110] Pandey AK, Ardekani BA, Kamarajan C, Zhang J, Chorlian DB, Byrne KN, Pandey G, Meyers JL, Kinreich S, Stimus A, Porjesz B (2018) Lower Prefrontal and Hippocampal Volume and Diffusion Tensor Imaging Differences Reflect Structural and Functional Abnormalities in Abstinent Individuals with Alcohol Use Disorder. Alcohol Clin Exp Res, 42(10), 1883–1896. https://doi.org/10.1111/acer.13854

[111] Fritz M, Klawonn AM, Zahr NM (2019) Neuroimaging in alcohol use disorder: From mouse to man. J Neurosci Res. https://doi.org/10.1002/jnr.24423

[112] Lee J, Im SJ, Lee SG, Stadlin A, Son JW, Shin CJ, Ju G, Lee SI, Kim S (2016) Volume of hippocampal subfields in patients with alcohol dependence. Psychiatry Res Neuroimaging, 258, 16–22. https://doi.org/10.1016/j.pscychresns.2016.10.009

[113] Oliveira AC, Pereira MC, Santana LN, Fernandes RM, Teixeira FB, Oliveira GB, Fernandes LM, Fontes-Junior EA, Prediger RD, Crespo-Lopez ME, Gomes-Leal W, Lima RR, Maia Cdo S (2015) Chronic ethanol exposure during adolescence through early adulthood in female rats induces emotional and memory deficits associated with morphological and molecular alterations in hippocampus. J Psychopharmacol, 29(6), 712–724. https://doi.org/10.1177/0269881115581960

[114] Staples MC, Mandyam CD (2016) Thinking after Drinking: Impaired Hippocampal-Dependent Cognition in Human Alcoholics and Animal Models of Alcohol Dependence. Front Psychiatry, 7, 162. https://doi.org/10.3389/fpsyt.2016.00162

[115] Akam T, Rodrigues-Vaz I, Marcelo I, Zhang X, Pereira M, Oliveira RF, Dayan P, Costa RM (2021) The Anterior Cingulate Cortex Predicts Future States to Mediate Model-Based Action Selection. Neuron, 109(1), 149–163 e147. https://doi.org/10.1016/j.neuron.2020.10.013

[116] Brockett AT, Tennyson SS, deBettencourt CA, Gaye F, Roesch MR (2020) Anterior cingulate cortex is necessary for adaptation of action plans. Proc Natl Acad Sci U S A, 117(11), 6196–6204. https://doi.org/10.1073/pnas.1919303117

[117] Lockwood PL, Wittmann MK (2018) Ventral anterior cingulate cortex and social decision-making. Neurosci Biobehav Rev, 92, 187–191. https://doi.org/10.1016/j.neubiorev.2018.05.030

[118] Rushworth MF, Walton ME, Kennerley SW, Bannerman DM (2004) Action sets and decisions in the medial frontal cortex. Trends Cogn Sci, 8(9), 410–417. https://doi.org/10.1016/j.tics.2004.07.009

[119] Kennerley SW, Walton ME, Behrens TE, Buckley MJ, Rushworth MF (2006) Optimal decision making and the anterior cingulate cortex. Nat Neurosci, 9(7), 940–947. https://doi.org/10.1038/nn1724

[120] Walton ME, Croxson PL, Behrens TE, Kennerley SW, Rushworth MF (2007) Adaptive decision making and value in the anterior cingulate cortex. Neuroimage, 36 Suppl 2, T142–154. https://doi.org/10.1016/j.neuroimage.2007.03.029

[121] Botvinick MM, Cohen JD, Carter CS (2004) Conflict monitoring and anterior cingulate cortex: an update. Trends Cogn Sci, 8(12), 539–546. https://doi.org/10.1016/j.tics.2004.10.003

[122] Carter CS, Braver TS, Barch DM, Botvinick MM, Noll D, Cohen JD (1998) Anterior cingulate cortex, error detection, and the online monitoring of performance. Science, 280(5364), 747–749. http://www.ncbi.nlm.nih.gov/entrez/query.fcgi?cmd=Retrieve&db=PubMed&dopt=Citation&list_uids=9563953

[123] van Veen V, Carter CS (2002) The anterior cingulate as a conflict monitor: fMRI and ERP studies. Physiol Behav, 77(4-5), 477–482. https://doi.org/10.1016/s0031-9384(02)00930-7

[124] Gehring WJ, Willoughby AR (2002) The medial frontal cortex and the rapid processing of monetary gains and losses. Science, 295(5563), 2279–2282. https://doi.org/10.1126/science.1066893

[125] Ma N, Liu Y, Li N, Wang CX, Zhang H, Jiang XF, Xu HS, Fu XM, Hu X, Zhang DR (2010) Addiction related alteration in resting-state brain connectivity. Neuroimage, 49(1), 738–744. https://doi.org/10.1016/j.neuroimage.2009.08.037

[126] Ojemann GA, Creutzfeldt O, Lettich E, Haglund MM (1988) Neuronal activity in human lateral temporal cortex related to short-term verbal memory, naming and reading. Brain, 111 (Pt 6)(6), 1383-1403. https://doi.org/10.1093/brain/111.6.1383

[127] Ojemann GA, Schoenfield-McNeill J, Corina D (2009) The roles of human lateral temporal cortical neuronal activity in recent verbal memory encoding. Cereb Cortex, 19(1), 197–205. https://doi.org/10.1093/cercor/bhn071

[128] Kable JW, Kan IP, Wilson A, Thompson-Schill SL, Chatterjee A (2005) Conceptual representations of action in the lateral temporal cortex. J Cogn Neurosci, 17(12), 1855–1870. https://doi.org/10.1162/089892905775008625

[129] Peelen MV, Romagno D, Caramazza A (2012) Independent representations of verbs and actions in left lateral temporal cortex. J Cogn Neurosci, 24(10), 2096–2107. https://doi.org/10.1162/jocn_a_00257

[130] Zhang S, Li CS (2014) Functional clustering of the human inferior parietal lobule by whole-brain connectivity mapping of resting-state functional magnetic resonance imaging signals. Brain Connect, 4(1), 53–69. https://doi.org/10.1089/brain.2013.0191

[131] Parsons OA, Nixon SJ (1993) Neurobehavioral sequelae of alcoholism. Neurol Clin, 11(1), 205–218. https://www.ncbi.nlm.nih.gov/pubmed/8441371

[132] Rao R, Topiwala A (2020) Alcohol use disorders and the brain. Addiction, 115(8), 1580–1589. https://doi.org/10.1111/add.15023

[133] Giancola PR, Moss HB (1998) Executive cognitive functioning in alcohol use disorders. Recent Dev Alcohol, 14, 227–251. https://doi.org/10.1007/0-306-47148-5_10

[134] Evert DL, Oscar-Berman M (1995) Alcohol-Related Cognitive Impairments: An Overview of How Alcoholism May Affect the Workings of the Brain. Alcohol Health Res World, 19(2), 89–96. https://www.ncbi.nlm.nih.gov/pubmed/31798082

[135] Oscar-Berman M, Shagrin B, Evert DL, Epstein C (1997) Impairments of brain and behavior: the neurological effects of alcohol. Alcohol Health Res World, 21(1), 65–75. https://www.ncbi.nlm.nih.gov/pubmed/15706764

[136] Fama R, Le Berre AP, Hardcastle C, Sassoon SA, Pfefferbaum A, Sullivan EV, Zahr NM (2019) Neurological, nutritional and alcohol consumption factors underlie cognitive and motor deficits in chronic alcoholism. Addict Biol, 24(2), 290–302. https://doi.org/10.1111/adb.12584

[137] Howard MO, Kivlahan D, Walker RD (1997) Cloninger’s tridimensional theory of personality and psychopathology: applications to substance use disorders. J Stud Alcohol, 58(1), 48–66. https://doi.org/10.15288/jsa.1997.58.48

[138] Conway KP, Compton W, Stinson FS, Grant BF (2006) Lifetime comorbidity of DSM-IV mood and anxiety disorders and specific drug use disorders: results from the National Epidemiologic Survey on Alcohol and Related Conditions. J Clin Psychiatry, 67(2), 247–257. https://doi.org/10.4088/jcp.v67n0211

[139] Kushner MG, Wall MM, Krueger RF, Sher KJ, Maurer E, Thuras P, Lee S (2012) Alcohol dependence is related to overall internalizing psychopathology load rather than to particular internalizing disorders: evidence from a national sample. Alcohol Clin Exp Res, 36(2), 325–331. https://doi.org/10.1111/j.1530-0277.2011.01604.x

[140] Nurnberger JI, Jr., Yang Z, Zang Y, Acion L, Bierut L, Bucholz K, Chan G, Dick DM, Edenberg HJ, Kramer J, Kuperman S, Rice JP, Schuckit M (2019) Development of Alcohol Use Disorder as a Function of Age, Severity, and Comorbidity with Externalizing and Internalizing Disorders in a Young Adult Cohort. J Psychiatr Brain Sci, 4. https://doi.org/10.20900/jpbs.20190016

[141] Meque I, Dachew BA, Maravilla JC, Salom C, Alati R (2019) Externalizing and internalizing symptoms in childhood and adolescence and the risk of alcohol use disorders in young adulthood: A meta-analysis of longitudinal studies. Aust N Z J Psychiatry, 53(10), 965–975. https://doi.org/10.1177/0004867419844308

[142] Hussong AM, Jones DJ, Stein GL, Baucom DH, Boeding S (2011) An internalizing pathway to alcohol use and disorder. Psychol Addict Behav, 25(3), 390–404. https://doi.org/10.1037/a0024519

[143] Weiss RD, Griffin ML, Mirin SM (1992) Drug abuse as self-medication for depression: an empirical study. Am J Drug Alcohol Abuse, 18(2), 121–129. https://doi.org/10.3109/00952999208992825

[144] Volkow ND (2004) The reality of comorbidity: depression and drug abuse. Biol Psychiatry, 56(10), 714–717. https://doi.org/10.1016/j.biopsych.2004.07.007

[145] Bottlender M, Soyka M (2005) Impact of different personality dimensions (NEO Five-Factor Inventory) on the outcome of alcohol-dependent patients 6 and 12 months after treatment. Psychiatry Res, 136(1), 61–67. https://doi.org/10.1016/j.psychres.2004.07.013

[146] Ribadier A, Varescon I (2019) Anxiety and depression in alcohol use disorder individuals: the role of personality and coping strategies. Subst Use Misuse, 54(9), 1475–1484. https://doi.org/10.1080/10826084.2019.1586950

[147] Dean SF, Fede SJ, Diazgranados N, Momenan R (2020) Addiction neurocircuitry and negative affect: A role for neuroticism in understanding amygdala connectivity and alcohol use disorder. Neurosci Lett, 722, 134773. https://doi.org/10.1016/j.neulet.2020.134773

[148] Griffith JW, Zinbarg RE, Craske MG, Mineka S, Rose RD, Waters AM, Sutton JM (2010) Neuroticism as a common dimension in the internalizing disorders. Psychol Med, 40(7), 1125–1136. https://doi.org/10.1017/S0033291709991449

[149] Kanner AD, Coyne JC, Schaefer C, Lazarus RS (1981) Comparison of two modes of stress measurement: daily hassles and uplifts versus major life events. J Behav Med, 4(1), 1–39. https://doi.org/10.1007/bf00844845

[150] Windle M, Windle RC (2015) A prospective study of stressful events, coping motives for drinking, and alcohol use among middle-aged adults. J Stud Alcohol Drugs, 76(3), 465–473. https://doi.org/10.15288/jsad.2015.76.465

[151] Bettis AH, Forehand R, McKee L, Dunbar JP, Watson KH, Compas BE (2016) Testing Specificity: Associations of Stress and Coping with Symptoms of Anxiety and Depression in Youth. J Child Fam Stud, 25(3), 949–958. https://doi.org/10.1007/s10826-015-0270-z

[152] Seiffge-Krenke I (2000) Causal links between stressful events, coping style, and adolescent symptomatology. J Adolesc, 23(6), 675–691. https://doi.org/10.1006/jado.2000.0352

[153] Edenberg HJ, Foroud T (2013) Genetics and alcoholism. Nat Rev Gastroenterol Hepatol, 10(8), 487–494. https://doi.org/10.1038/nrgastro.2013.86

[154] Verhulst B, Neale MC, Kendler KS (2015) The heritability of alcohol use disorders: a meta-analysis of twin and adoption studies. Psychol Med, 45(5), 1061–1072. https://doi.org/10.1017/S0033291714002165

[155] Friedel E, Kaminski J, Ripke S (2021) Heritability of Alcohol Use Disorder: Evidence from Twin Studies and Genome-Wide Association Studies. In: Textbook of Addiction Treatment, Springer, pp. 21–33. https://link.springer.com/chapter/10.1007/978-3-030-36391-8_3

[156] Johnson EC, Sanchez-Roige S, Acion L, Adams MJ, Bucholz KK, Chan G, Chao MJ, Chorlian DB, Dick DM, Edenberg HJ, Foroud T, Hayward C, Heron J, Hesselbrock V, Hickman M, Kendler KS, Kinreich S, Kramer J, Kuo SI, Kuperman S, Lai D, McIntosh AM, Meyers JL, Plawecki MH, Porjesz B, Porteous D, Schuckit MA, Su J, Zang Y, Palmer AA, Agrawal A, Clarke TK, Edwards AC (2021) Polygenic contributions to alcohol use and alcohol use disorders across population-based and clinically ascertained samples. Psychol Med, 51(7), 1147–1156. https://doi.org/10.1017/S0033291719004045

[157] Meyers JL, Chorlian DB, Johnson EC, Pandey AK, Kamarajan C, Salvatore JE, Aliev F, Subbie-Saenz de Viteri S, Zhang J, Chao M, Kapoor M, Hesselbrock V, Kramer J, Kuperman S, Nurnberger J, Tischfield J, Goate A, Foroud T, Dick DM, Edenberg HJ, Agrawal A, Porjesz B (2019) Association of Polygenic Liability for Alcohol Dependence and EEG Connectivity in Adolescence and Young Adulthood. Brain Sci, 9(10). https://doi.org/10.3390/brainsci9100280

[158] Clarke TK, Smith AH, Gelernter J, Kranzler HR, Farrer LA, Hall LS, Fernandez-Pujals AM, MacIntyre DJ, Smith BH, Hocking LJ, Padmanabhan S, Hayward C, Thomson PA, Porteous DJ, Deary IJ, McIntosh AM (2016) Polygenic risk for alcohol dependence associates with alcohol consumption, cognitive function and social deprivation in a population-based cohort. Addict Biol, 21(2), 469–480. https://doi.org/10.1111/adb.12245

[159] Hatoum AS, Johnson EC, Baranger DAA, Paul SE, Agrawal A, Bogdan R (2021) Polygenic risk scores for alcohol involvement relate to brain structure in substance-naive children: Results from the ABCD study. *Genes Brain Behav*, e12756. https://doi.org/10.1111/gbb.12756

[160] Roy A, Svensson FP, Mazeh A, Kocsis B (2017) Prefrontal-hippocampal coupling by theta rhythm and by 2-5 Hz oscillation in the delta band: The role of the nucleus reuniens of the thalamus. Brain Struct Funct, 222(6), 2819–2830. https://doi.org/10.1007/s00429-017-1374-6

[161] Ketz NA, Jensen O, O’Reilly RC (2015) Thalamic pathways underlying prefrontal cortex-medial temporal lobe oscillatory interactions. Trends Neurosci, 38(1), 3–12. https://doi.org/10.1016/j.tins.2014.09.007

[162] Yamamoto H, Kubota S, Shimizu FA, Hirano-Iwata A, Niwano M (2018) Effective Subnetwork Topology for Synchronizing Interconnected Networks of Coupled Phase Oscillators. Front Comput Neurosci, 12, 17. https://doi.org/10.3389/fncom.2018.00017

[163] Babapoor-Farrokhran S, Vinck M, Womelsdorf T, Everling S (2017) Theta and beta synchrony coordinate frontal eye fields and anterior cingulate cortex during sensorimotor mapping. Nat Commun, 8, 13967. https://doi.org/10.1038/ncomms13967

[164] Axmacher N, Henseler MM, Jensen O, Weinreich I, Elger CE, Fell J (2010) Cross-frequency coupling supports multi-item working memory in the human hippocampus. Proc Natl Acad Sci U S A, 107(7), 3228–3233. https://doi.org/10.1073/pnas.0911531107

[165] Brincat SL, Donoghue JA, Mahnke MK, Kornblith S, Lundqvist M, Miller EK (2021) Interhemispheric transfer of working memories. Neuron, 109(6), 1055–1066 e1054. https://doi.org/10.1016/j.neuron.2021.01.016

[166] Graham K, Bernards S, Knibbe R, Kairouz S, Kuntsche S, Wilsnack SC, Greenfield TK, Dietze P, Obot I, Gmel G (2011) Alcohol-related negative consequences among drinkers around the world. Addiction, 106(8), 1391–1405. https://doi.org/10.1111/j.1360-0443.2011.03425.x

[167] Reid MC, Fiellin DA, O’Connor PG (1999) Hazardous and harmful alcohol consumption in primary care. Arch Intern Med, 159(15), 1681–1689. https://doi.org/10.1001/archinte.159.15.1681

[168] Tuominen L (2014) Neurobiological correlates of personality traits: A Study on harm avoidance and neuroticism. University of Turku, Turku. http://www.antoniocasella.eu/archipsy/Tuominen_2014.pdf

