## SupplementaryMaterial for "Predicting alcohol-related memory problems in older adults: A machine learning study with multi-domain features"

### 1. Material and Methods

#### 1.1. Sample Description

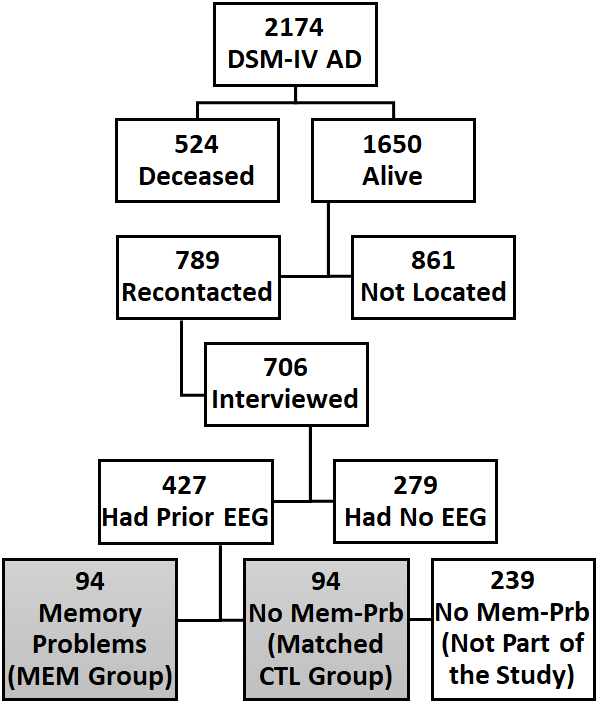

Fig. S1. Flowchart showing the selection process for the study sample. *Abbreviation:* AD **–** Alcohol dependence; MEM **–** Memory Group; CTL **–** Comparison Group; Mem-Prb **–** Memory Problems.

The data were collected from six collection centers of the COGA study (SUNY Brooklyn, University of Connecticut, Indiana University, Washington University in St. Louis, University of Iowa, and University of California, San Diego). A phone interview was conducted to assess current functional and health status, including alcohol use and memory problems. All living participants provided informed consent in compliance with their local IRBs. The flow chart for the sample selection for the current study is illustrated in **Fig. S1**. Initially, we identified 2174 older COGA participants who: (i) met the criteria for lifetime DSM-IV alcohol dependence based on the Semi-Structured Assessment for the Genetics of Alcohol (SSAGA) [1,2] based on prior interviews during earlier phases of COGA, (ii) were born before 1967 (thus to be aged 50 or older at the assessment), and (iii) had DNA collected. Circumstances dictated that the study be conducted over a 12-month period starting in January 2017. Out of the 2174 eligible participants, 524 were deceased, and of those that were alive, 789 were contacted (861 were not contacted for various reasons). Out of the 789 who were re-contacted, 706 were administered a brief telephone interview for a recent follow-up study. The present study sample was selected from 427 individuals who had prior EEG recording during the earlier phases of COGA over more than 18 years ago (Mean=18.53; SD=3.86). Among those with EEG, 94 individuals, who had endorsed having alcohol-related memory problems during the past 5 years and/or 10 years (self-report) during the recent follow up telephone interview, formed the experimental group with memory problems (*Memory group*). The matched comparison control group (*Control group*, N=94) was selected from those with EEG but without having any reported memory problems by matching based on several parameters: age at EEG recording, sex, ethnicity, and past alcohol use pattern during the latest SSAGA interview.

#### 1.2. Recent Follow-up Telephone Interview

The items of the recent follow-up telephone interview covered the following categories: (a) marital status, (b) living arrangements, (c) education, (d) employment, (e) physical health, (f) mental health; (g) alcohol use and related health consequences (quantity/frequency, memory, blackouts, etc.); and (h) willingness to participate again in a future assessment. Alcohol-related questions consisted of prior 5-year and 10-year alcohol problems that included alcohol-related blackouts, difficulties maintaining drinking limits, spending much significant time using or recovering from its effects, interpersonal or work/school problems, use in hazardous situations, problems cutting back or stopping use, alcohol-related health impairment, and signs of alcohol withdrawal. Items about the quantity/frequency of alcohol use were queried for the past 12 months before the interview. Self-ratings of current physical and mental health were each scored on a 4-point scale from excellent to poor, and self-rating of memory on a 3-point scale of better, the same, or worse compared to other people their age. Additional details of the interview items and related data are available from our previous publications [3,4]. The list of variables from the follow-up interview schedule (N=12) are listed below in **Table S1**.

Table S1: The list of variables from the follow-up interview schedule (N=12) included in the random forest classification model.

| Feature | Description | Instrument / Source |
| --- | --- | --- |
| PhyHealth | Current state of physical health | Followup interview schedule |
| MenHealth | Current state of mental health | Followup interview schedule |
| DaysLastDrk | Days since last full standard drink | Followup interview schedule |
| WeeksDrk | in the past 12 months, number of weeks with alcohol consumption (max=52 weeks) | Followup interview schedule |
| Drk24Hr | In the past 12 months, the largest number of drinks in a 24-hour period | Followup interview schedule |
| DaysAbst | In the last 12 months, the longest period without drinking (in number of days) | Followup interview schedule |
| AlcExp5yrs | Number of alcohol-related negative experiences or symptoms in the last 5 years (max=6) | Followup interview schedule |
| AlcWthSx5yrs | Number of alcohol withdrawals symptoms in the last 5 years (max=5) | Followup interview schedule |
| AlcHlthProb5yrs | Number of health problems in the last 5 years (max=5) | Followup interview schedule |
| AlcExp10yrs | Number of alcohol-related negative experiences or symptoms in the last 10 years (max=6) | Followup interview schedule |
| AlcWthSx10yrs | Number of alcohol withdrawals symptoms in the last 10 years (max=5) | Followup interview schedule |
| AlcHlthProb10yrs | Number of health problems in the last10 years (max=5) | Followup interview schedule |

#### 1.3. EEG Data Acquisition and Preprocessing

Prior to the EEG recording, participants were asked to have abstained from alcohol for a minimum of 5 days. Individuals were excluded from the recording if they reported any of the following: (1) recent alcohol use in the past 5 days (i.e., positive breath-analyzer test); (2) hepatic encephalopathy/cirrhosis of the liver; (3) significant history of head injury, seizures or neurosurgery; (4) uncorrected sensory deficits; (5) taking medication known to influence brain functioning; and (6) other acute/chronic medical/neurological illnesses that affect brain function (multiple sclerosis, meningitis, encephalitis, neurodegenerative diseases, stroke, traumatic brain injury, brain tumor, etc.). Participants were seated comfortably in a dimly lit sound-attenuated, temperature-regulated booth (Industrial Acoustics, Bronx, NY, USA). EEG was recorded during the awake, eyes-closed resting state for 4.25 minutes, either using a MASSCOMP 5550 system (Concurrent Computer Corporation, Duluth, GA, USA) at the sampling rate of 256 Hz with bandpass between 0.02–50 Hz or using a Neuroscan system (Version 4.1) (Compumedics Limited, Charlotte, NC, USA) at a sampling rate of either 500 Hz or 512 Hz with bandpass between 0.02–100.0 Hz. EEG signals were recorded from the scalp were recorded using an electrode cap (Electro-Cap International, Eaton, OH, USA) with a 19-channel montage of the 10–20 international system [5-7], and were amplified 10,000 times by either Sensorium (Charlotte, VT, USA) EPA-2 or Neuroscan amplifiers. The reference electrode was fixed on the nose tip, and a forehead electrode served as the ground. The electrooculogram (EOG) was recorded by a supraorbital vertical electrode and by a horizontal electrode on the external canthus of the left eye. Electrode impedances were maintained below 5 kΩ. EEG acquisition protocol was identical across all six collection sites of COGA [8,9].

As described in our previous work on EEG source functional connectivity [10], preprocessing was performed using custom scripts in Matlab (The MathWorks, Inc., Natick, MA) at two levels: (a) on the entire continuous EEG recording and (b) on each of the segmented epochs. The following steps were performed on the entire continuous EEG trace of the recording: (i) data points were resampled to 256 Hz for harmonizing different sampling rates; (ii) bandpass filtering at 0.05–50 Hz to keep only the frequency range of interest; (iii) waveforms were "detrended" to remove low frequency components resulting in a near linear upward/downward trending deviation; and (iv) "de-meaning" was done by subtracting the gross mean of the entire EEG trace from each data point in order to align the waveforms close to the zero-amplitude baseline. Then, the continuous EEG data were segmented into epochs of 2000 ms. The next batch of preprocessing steps was performed on each of the epochs: (i) detrending; (ii) baseline alignment by subtracting epoch mean from each data point; (iii) interpolation of missing data or "flat" channels by computing the mean of surrounding nearest channels; (iv) removal of epochs with DC shift/drift involving voltage steps higher than 75 mV between any two adjacent sampling points; and (v) removal of possible EOG contaminated epochs if any data point was beyond the threshold of ±100 μV or if the difference between lowest and highest amplitude within the epoch was 200 μV. Artifact free 30 artifact free random epochs were selected randomly for the functional connectivity analysis to keep a uniform minimum number of epochs across subjects.

#### 1.4. EEG Functional Connectivity Analysis using eLORETA

The eLORETA software [11,12] includes several different methods to analyze EEG source data (current density) in 3-dimensional space with 6239 voxels at 5×5×5 mm spatial resolution. The functional connectivity across the default mode network regions (**Fig. S2**) were computed using the EEG source data based on lagged linear connectivity (LLC), which is less susceptible to volume conduction artifacts of the scalp-recorded EEG signals (Pascual-Marqui et al, 2007). The procedure adapted in the study has been detailed in our previous publications [10,13]. Lagged phase synchronization is a measure of similarity (a corrected phase synchrony value) between signals in the frequency domain based on normalized Fourier transforms [12], representing the strength of connectivity between two signals by subtracting the instantaneous zero-lag (non-physiological) contribution from the total connectivity to retain only the physiological connectivity between true cortical sources [12,14]. As explained by Canuet et al. [14], while the classical coherence or connectivity measure, which contains both real and imaginary component, is represented as:

(i)
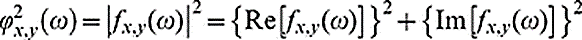
, in which
(ii)
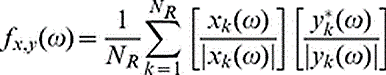

On the other hand, the Lagged phase synchronization, which statistically partials out the instantaneous component of the total connectivity, is defined as:

(iii)
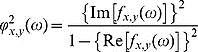

In the equations (i) and (iii), *Re* and *Im* denote the real and imaginary parts of a complex number. In the equation (ii), xk(ω) and yk(ω) denote the discrete Fourier transforms of x and y at frequency ω for the k-th EEG segment or epoch (k=1…NR), in which NR is the number of epochs. Thus, lagged phase synchronization, which is devoid of the instantaneous component of the classical coherence measure, is a powerful measure to elicit uncontaminated functional connectivity between the signals of interest.

#### 1.5. Functional Connectivity Across the Default Mode Network

The default mode network regions analyzed in the study are illustrated in **Fig. S2**. Each seed region contained the voxels within a 10 mm radius from the peak/centroid point of the region. The ROI-to-ROI connectivity [15], the most commonly used method to derive functional connectivity across brain regions [16], was computed using EEG based functional connectivity was computed using the exact Low Resolution Electric Tomography software (eLORETA) software [11,12] for the for the custom frequency bands: delta (1–3 Hz), theta (4–7 Hz), alpha (8–12 Hz), beta (13–29 Hz), and gamma (30–40 Hz).

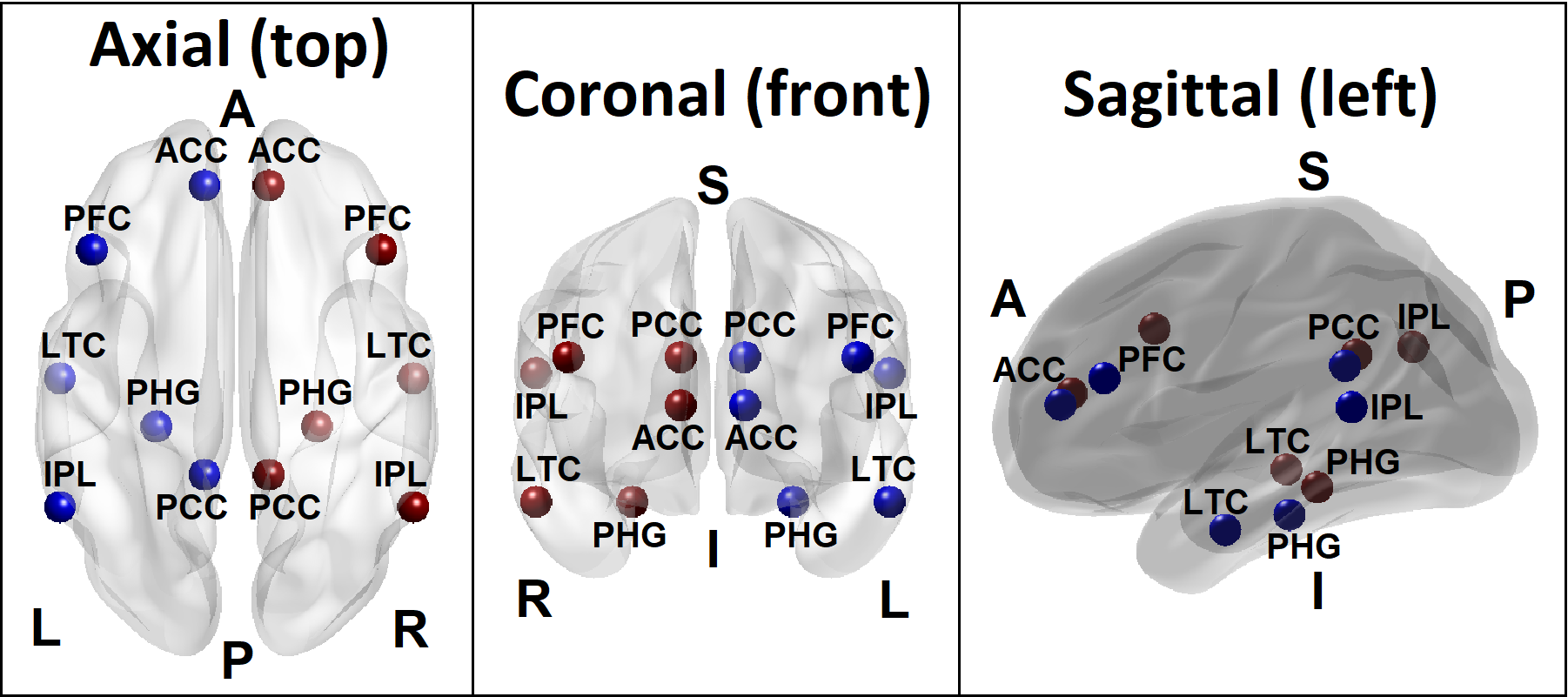

Fig. S2. The Default Mode Network nodes included for the computation of functional connectivity: Six bilateral default mode network nodes from which were functional connectivity was calculated include: prefrontal cortex (PFC), anterior cingulate cortex (ACC), posterior cingulate cortex (PCC), hippocampal formation (HCF), inferior parietal lobule (IPL), and lateral temporal cortex (LTC) in each hemisphere. [Views: Top view–left panel; Front view–middle panel; Side view–right panel. Direction: A–Anterior; P–Posterior; L–Left; R–Right; S–Superior; I–Inferior]

#### 1.6. Assessment of Temperament, Personality, and Alcohol Experience

The seven questionnaires from which behavioral data included for the current study were: (i) Sensation Seeking Scale (SSS) Form-V [17], a 40-item self-report measure consisting of four subscales that include thrill and adventure seeking (TAS), experience seeking (ES), disinhibition (Dis), and boredom susceptibility (BS); (ii) Tri-Dimensional Personality Questionnaire (TPQ) [18], a 100-item self-report measure to assess novelty seeking (NS), harm avoidance (HA), and reward dependence (RD); (iii) Daily Hassles and Uplifts (DHU) [19], a 53-item scale to measure cumulative indices of hassles (HSL) and uplifts (UPL); (iv) NEO Five Factor Inventory (NEO) [20], a 60-item self-report questionnaire that measures five domains of adult personality, including such as neuroticism (N), extroversion (E), openness to experience (O), agreeableness (A), and conscientiousness (C); (v) Perceived Social Support (PSS) [21] to measure perceived support from family (PSS-FA) and friends (PSS-FR); (vi) Alcohol Expectancy Questionnaire Adult (AEQ) [22], a 120-item self-report form to measure the respondent's beliefs about the effects of alcohol on global positive changes (GPC), enhanced sexuality (ES), physical and social pleasure (PSP), increased social assertiveness (ISA), arousal and aggression (AA), and relaxation and tension reduction (RTR); and (vii) Self Rating of Response to Ethanol (SRE) [23], a self-report instrument to measure subjective and actual effects of drinking for the first 5 drinking episodes (SRE-5drk), first 3 months of regular drinking (SRE-3mon), and period of heaviest drinking (SRE-Hvy). These variables are listed in **Table S2** below.

Table S2: The list of variables (N=27) for the personality and life experiences questionnaires.

| Feature | Instrument | Details |
| --- | --- | --- |
| SSV_DIS | Sensation Seeking Scale Form-V (SSV) | SSV score for Disinhibition (DIS) subscale |
| SSV_BS | Sensation Seeking Scale Form-V (SSV) | SSV score for Boredom Susceptibility (BS) subscale |
| SSV_TAS | Sensation Seeking Scale Form-V (SSV) | SSV score for Thrill and Adventure Seeking (TAS) subscale |
| SSV_ES | Sensation Seeking Scale Form-V (SSV) | SSV score for Experience Seeking (ES) subscale |
| SSV_TOT | Sensation Seeking Scale Form-V (SSV) | SSV Total score |
| TPQ_NS | Tri-Dimensional Personality Questionnaire (TPQ) | TPQ score for Novelty Seeking (NS) category |
| TPQ_HA | Tri-Dimensional Personality Questionnaire (TPQ) | TPQ score for Harm Avoidance (HA) category |
| TPQ_RD | Tri-Dimensional Personality Questionnaire (TPQ) | TPQ score for Reward Dependence (RD) category |
| AEQ_GPC | Alcohol Expectancy Questionnaire Adult (AEQ) | AEQ score for Global Positive Changes (GPC) subscale |
| AEQ_ES | Alcohol Expectancy Questionnaire Adult (AEQ) | AEQ score for Enhanced Sexuality (ES) subscale |
| AEQ_PSP | Alcohol Expectancy Questionnaire Adult (AEQ) | AEQ score for Physical and Social Pleasure (PSP) subscale |
| AEQ_ISA | Alcohol Expectancy Questionnaire Adult (AEQ) | AEQ score for Increased Social Assertiveness (ISA) subscale |
| AEQ_RTR | Alcohol Expectancy Questionnaire Adult (AEQ) | AEQ score for Relaxation and Tension Reduction (RTR) subscale |
| AEQ_AP | Alcohol Expectancy Questionnaire Adult (AEQ) | AEQ score for Arousal and Aggression (AA) subscale |
| DHU_HSL | Daily Hassles and Uplifts (DHU) | DHU score for Hassles (HSL) category |
| DHU_UPL | Daily Hassles and Uplifts (DHU) | DHU score for Uplifts (UPL) category |
| NEO_N | NEO Five Factor Inventory (NEO) | NEO score for Neuroticism (N) category |
| NEO_E | NEO Five Factor Inventory (NEO) | NEO score for Extroversion (E) category |
| NEO_O | NEO Five Factor Inventory (NEO) | NEO score for Openness to experience (O) category |
| NEO_A | NEO Five Factor Inventory (NEO) | NEO score for Agreeableness (A) category |
| NEO_C | NEO Five Factor Inventory (NEO) | NEO score for Conscientiousness (C) category |
| SRE_5drk | Self Rating of Response to Ethanol (SRE) | SRE score for the effects of drinking during the first 5 drinking occasions (5drk) |
| SRE_Hvy | Self Rating of Response to Ethanol (SRE) | SRE score for the effects of drinking during the heviest drinking period (Hvy) |
| SRE_3mon | Self Rating of Response to Ethanol (SRE) | SRE score for the effects of drinking during the last 3 months (3mon) |
| SRE_Tot | Self Rating of Response to Ethanol (SRE) | SRE total score |
| PSS_Fam | Perceived Social Support (PSS) | PSS score for the Family (Fam) scale |
| PSS_Frs | Perceived Social Support (PSS) | PSS score for the Friends (Frs) scale |

#### 1.7. Genomic Data and Polygenic Risk Scores (PRS)

Genotyping of the COGA data was conducted across different phases of data collection, and genotyped at multiple sites, including (i) Center for Inherited Disease Research using the Illumina HumanHap1M array [24]; (ii) Genome Technology Access Center at Washington University School of Medicine using the Illumina OmniExpress [25]; and (iii) Rutgers University using the Affymetrix Smokescreen array [26]. Data were imputed to 1000 Genomes (Phase 3, version 5) using SHAPEIT [27] and then Minimac3 [28]. Genotyping arrays were imputed separately due to different variant contents on each array. Prior to imputation, variants with missing rates > 5%, MAF < 3% and HWE p values <0.0001 were excluded. Following imputation, genotype probabilities ≥ 0.90 were changed to genotypes. Mendelian errors in the imputed SNPs were reviewed and resolved as described previously [29,30]. SNPs with an imputation information score < 0.30 or MAF < 0.03 were excluded from subsequent analysis.

The phenotypes for which PRS calculations were performed were the following: (i) AUD diagnosis based on ICD-9 or ICD-10 codes [31], (ii) AUDIT-C scores [31], and (iii) maximum habitual alcohol intake [32] from the Million Veteran Program (MVP), and DSM-IV alcohol dependence [33] from the Psychiatric Genomic Consortium (PGC). Each of these GWAS had samples from both European Ancestry (EA) and African Ancestry (AA). However, PRS of neurocognitive phenotypes, specifically memory functions, was not included in the study due to a lack of availability of multi-ethnic PRS data. The PRS-CSx [34-37], used in the current study, is an extension of PRS-CS [34], has been primarily implemented for cross-ethnic polygenic prediction. This method integrates summary statistics of GWAS and external LD reference panels from multiple populations to improve cross-population polygenic prediction. For the current study, we used only the SNPs that were common to both European and African ancestries. We also limited the SNPs for score creation to HapMap3 SNPs that overlapped between the original GWAS summary statistics, the LD reference panels (1000 Genomes Phase III European and African subsamples), and the target samples for score creation. PRS were converted to Z-scores for interpretability.

#### 1.8. Feature selection of EEG Functional Connectivity variables

In the current study, feature selection method was used as a first stage to reduce irrelevant and redundant variables which may otherwise add noise to the predictive models [38-40]. Advantages of feature selection include a better understanding of the data quality, reducing computation requirements, reducing the effect of the curse of dimensionality (problems, such as sparsity, related to high-dimensional datasets), and also improving the predictor performance [39]. In the current study, we applied binomial lasso regression as the feature selection method as implemented in R-package 'glmnet' [41], in which generalized linear (used here) and similar models are fitted via penalized maximum likelihood and the regularization path is computed for the lasso (used here) or elastic net penalty at a grid of values for the regularization parameter *lambda*. At the first step, a binomial logistic regression model of classification type involving model fit criteria (e.g., 10% yield). In the second step, beta coefficients for a specific lambda criterion (“lambda.min” or “lambda.1se”) are derived and only the features with “non-zero” coefficients (i.e., the features with signals or classificatory power) are selected. Using this method, a set of EEG functional connectivity variables with significant predictive values to discriminate *Memory* group from the *Control* group. The method adapted in the current analysis is based on the Lasso method as implemented in Fonti and Belitser [42], in which the model included the maximum output features "pmax" was set to 10% (i.e., 330 ÷ 10% = 33 variables). The 10-fold cross-validation, coupled with lambda thresholding at 1 SE (λ_1se_), was used to extract the final set of key variables, while the area under the curve (AUC) was used to assess the classification performance of selected features.

#### 1.9. Random Forests classification model and parameters

Random Forests, an efficient predictive algorithm, was devised by Breiman [43]. The classifier algorithm consists of a collection of tree-structured classifiers where each tree casts a unit vote for a class/group for each set of predictor variables. A growing number of studies in computational biology are using Random Forests because of several advantages of the method. According to Qi [44], the Random Forests method is not only nonparametric but is interpretable and efficient. Further, the Random Forests method can be applied to data with a small sample size, multi-dimensional variables, and multiple layers/levels without compromising its prediction accuracy [44]. In a large-scale benchmark experiment, the Random Forests algorithm was found to perform better than logistic regression in terms of prediction accuracy [45]. The two main parameters of the Random Forests algorithm are the number of trees in the ensemble and the number of variables randomly selected for the splitting decision at each node. Two levels of randomness are used by the Random Forests to construct the ensemble of trees: first, the model trains itself using training data for creating each tree based on bootstrap aggregating (bagging). At the second level, the algorithm randomly selects a subset of features to split at each node while growing a decision tree for group classification. To maximize the classification accuracy (by reducing the errors or impurity), only a single best feature (variable) among a random subset of features is selected at each internal node. This process is recursively repeated until one of the three conditions is met: (i) the tree has either reached a specified depth (i.e., number of layers of splits between the original data and the data at the bottom of the decision tree), (ii) the number of samples in a node becomes lower than the set threshold, and (iii) when all the samples are grouped into the same category [46]. Some of the important concepts and parameters of Random Forests classification method are listed in *Box S1*.

**Box S1:** Concepts and parameters used in Random Forest classification method

- **Trees:** Decision trees whose results are aggregated into one final result for classifying the factors or outcomes. Each tree is constructed based on a random (bootstrapped) subsample of the observations.
- **Node:** A point in a tree, where a split occurs as a result of a ’test’ on an attribute leading to binary outcomes (e.g., whether a coin flip results in head or tail). A binary split at a node partitions the data from the parent node into two daughter nodes.
- **Branch:** The outcome of the test resulting in a split or two branches in a classification tree.
- **Leaf:** A terminal node that has no children or branches.
- **Random Forest ensemble:** Aggregation of individual decision trees in order to combine predictions (votes) from each tree. The class/group/outcome with most votes becomes the Random Forests model’s prediction.
- **Bagging:** It’s the short form of ’bootstrap aggregating’, which is a method to improve classification by combining classifications of randomly generated training sets.
- **Out of bag (OOB) estimate:** The observations that are not part of the bootstrap subsample are referred to as out-of-bag (OOB) observations. The OOB error refers to the classification error based on this subsample and serves as a validation of Random Forest model accuracy.
- **Gini (mean) decrease:** It represents the importance of a specific feature/predictor/variable (Vi) for the classification or prediction. It’s the mean decrease in node impurity (classification error) of Vi. A higher Gini decrease indicates higher variable importance for Vi.
- **Accuracy decrease:** Mean decrease in prediction accuracy after Vi is not taken into account.
- **Mean minimal depth:** It refers to the number of nodes along the shortest path from the root node down to the nearest leaf node. Smaller depth for the Vi indicates its higher importance.
- **Mtry:** A preset number of features/variables/predictors randomly selected (from the entire list) for splitting at each node in the construction of each decision tree.
- **ntree:** A preset total number of trees to grow for a given model. Larger ‘ntree’ normally produce more stable models and more reliable predictions.
- **Number of nodes:** Total number of nodes that use Vi for splitting (it is usually equal to number of trees if trees are shallow).
- **Times a root:** Total number of trees in which Vi is used for splitting the root node (i.e., the whole sample is divided into two based on the value of Vi).
- **P-value:** Probability value of hypothesis testing based on a one-sided binomial test that indicates whether the observed number of successes (number of nodes in which Vi was used for splitting) exceeds the theoretical number of successes if they were random.

To compute prediction error and classification accuracy, we used the Out-of-Bag (OOB) error estimate, which represents the classification error obtained from the out-of-bag sample (about one-third of the total sample) that was not part of the bootstrap sample (about two-third of the total sample) used in growing the forests. In the Random Forests model, cross-validation in a separate test sample is not required, as it is estimated internally in the algorithm [47]. During each iteration of constructing a decision tree, about two-thirds of the bootstrap sample from the training data is used, and about one-third of the sample is left out during each bootstrap process, which is called the out-of-bag (OOB) sample. The classification error calculated from this sample is called the OOB error score. The aggregate of OOB scores from all decision trees will provides the ensemble OOB error rate (i.e., classification error) as well as the Random Forests model accuracy rate for the Random Forests model. Thus, the OOB score provides validation for the Random Forests model. The Random Forests classification model included 29 default mode network connections from feature selection (see Results section), 27 variables on temperament, personality, and life experiences, 12 variables on health and alcohol-related problems, and 4 PRS scores on alcohol phenotypes as features, while the group status (*Memory* vs. *Control* group) served as the outcome variable. In the model, the maximum number of trees ‘ntree’ was set at 500. The optimal number of features analyzed at each node ('Mtry') was estimated to be 9 (using the ‘tuneRF’ function) and was used in the classifier algorithm. The final list of variables that significantly contributed to the classification was tabulated, and 3-dimensional connectivity maps of top significant default mode network connections within a brain anatomical template were created using custom Matlab scripts.

### 2. Results

#### 2.1. Feature Selection of EEG Functional Connectivity variables

The feature selection procedure identified a total of 29 functional connectivity variables from multiple frequency bands connecting across the twelve default mode network seeds (see **Table S3** below**)**. These connections are: Delta – 12 connections (1–5, 1–6, 1–12, 2–12, 2–5, 3–5, 3–7, 4–8, 5–6, 6–11, 7–11, 8–12), Theta – 6 connections (2–5, 2–11, 4–10, 4–6, 6–10, 9–11), Alpha – 4 connections (2–5, 2–11, 7–10, 7–12), Beta – 5 (1–4, 2–10, 3–7, 4–9, 5–12), and Gamma – 2 variables (2–10, 4–12).

Table S3: The list of functional connectivity variables (N=29) that were identified by the feature selection method and included in the random forest classification analysis.

| Feature | Frequency | Node 1 | Node 2 |
| --- | --- | --- | --- |
| FC_De_1_5 | Delta | Left posterior cingulate cortex | Left inferior parietal lobule |
| FC_De_1_6 | Delta | Left posterior cingulate cortex | Right inferior parietal lobule |
| FC_De_1_12 | Delta | Left posterior cingulate cortex | Right parahippocampal gyrus |
| FC_De_2_5 | Delta | Right posterior cingulate cortex | Left inferior parietal lobule |
| FC_De_2_12 | Delta | Right posterior cingulate cortex | Right parahippocampal gyrus |
| FC_De_3_5 | Delta | Left anterior cingulate cortex | Left inferior parietal lobule |
| FC_De_3_7 | Delta | Left anterior cingulate cortex | Left prefrontal cortex |
| FC_De_4_8 | Delta | Right anterior cingulate cortex | Right prefrontal cortex |
| FC_De_5_6 | Delta | Left inferior parietal lobule | Right inferior parietal lobule |
| FC_De_6_11 | Delta | Right inferior parietal lobule | Left parahippocampal gyrus |
| FC_De_7_11 | Delta | Left prefrontal cortex | Left parahippocampal gyrus |
| FC_De_8_12 | Delta | Right prefrontal cortex | Right parahippocampal gyrus |
| FC_Th_2_5 | Theta | Right posterior cingulate cortex | Left inferior parietal lobule |
| FC_Th_2_11 | Theta | Right posterior cingulate cortex | Left parahippocampal gyrus |
| FC_Th_4_6 | Theta | Right anterior cingulate cortex | Right inferior parietal lobule |
| FC_Th_4_10 | Theta | Right anterior cingulate cortex | Right lateral temporal cortex |
| FC_Th_6_10 | Theta | Right inferior parietal lobule | Right lateral temporal cortex |
| FC_Th_9_11 | Theta | Left lateral temporal cortex | Left parahippocampal gyrus |
| FC_Al_2_5 | Alpha | Right posterior cingulate cortex | Left inferior parietal lobule |
| FC_Al_2_11 | Alpha | Right posterior cingulate cortex | Left parahippocampal gyrus |
| FC_Al_7_10 | Alpha | Left prefrontal cortex | Right lateral temporal cortex |
| FC_Al_7_12 | Alpha | Left prefrontal cortex | Right parahippocampal gyrus |
| FC_Be_1_4 | Beta | Left posterior cingulate cortex | Right anterior cingulate cortex |
| FC_Be_2_10 | Beta | Right posterior cingulate cortex | Right lateral temporal cortex |
| FC_Be_3_7 | Beta | Left anterior cingulate cortex | Left prefrontal cortex |
| FC_Be_4_9 | Beta | Right anterior cingulate cortex | Left lateral temporal cortex |
| FC_Be_5_12 | Beta | Left inferior parietal lobule | Right parahippocampal gyrus |
| FC_Ga_2_10 | Gamma | Right posterior cingulate cortex | Right lateral temporal cortex |
| FC_Ga_4_12 | Gamma | Right anterior cingulate cortex | Right parahippocampal gyrus |

**Abbreviations:** FC­–Functional Connectivity; De–Delta; Th–Theta; Al–Alpha; Be–Beta; Ga–Gamma; Numbers in functional connectivity variables: 1-12 of the default mode network.

#### 2.2. Random Forests Classification Accuracy

The overall prediction accuracy of the Random Forests model to classify *Memory* and *Control* individuals using FC, PRS, behavioral and clinical predictors, as estimated by the area under the ROC curve, was 88.29% (**Fig. S3**). The confusion matrix of the Random Forests model showed that the accuracy rate for *Memory* and *Control* groups were 72.34% and 90.43%, respectively, with accuracy rate 81.39%, sensitivity 88.31%, specificity 76.58%, positive predictive value 72.34% and negative predictive value 90.43%. In other words, 68 individuals in the *Memory* group and 85 individuals in the *Control* group (out of 94 in each group) were correctly identified by the Random Forests classification algorithm.

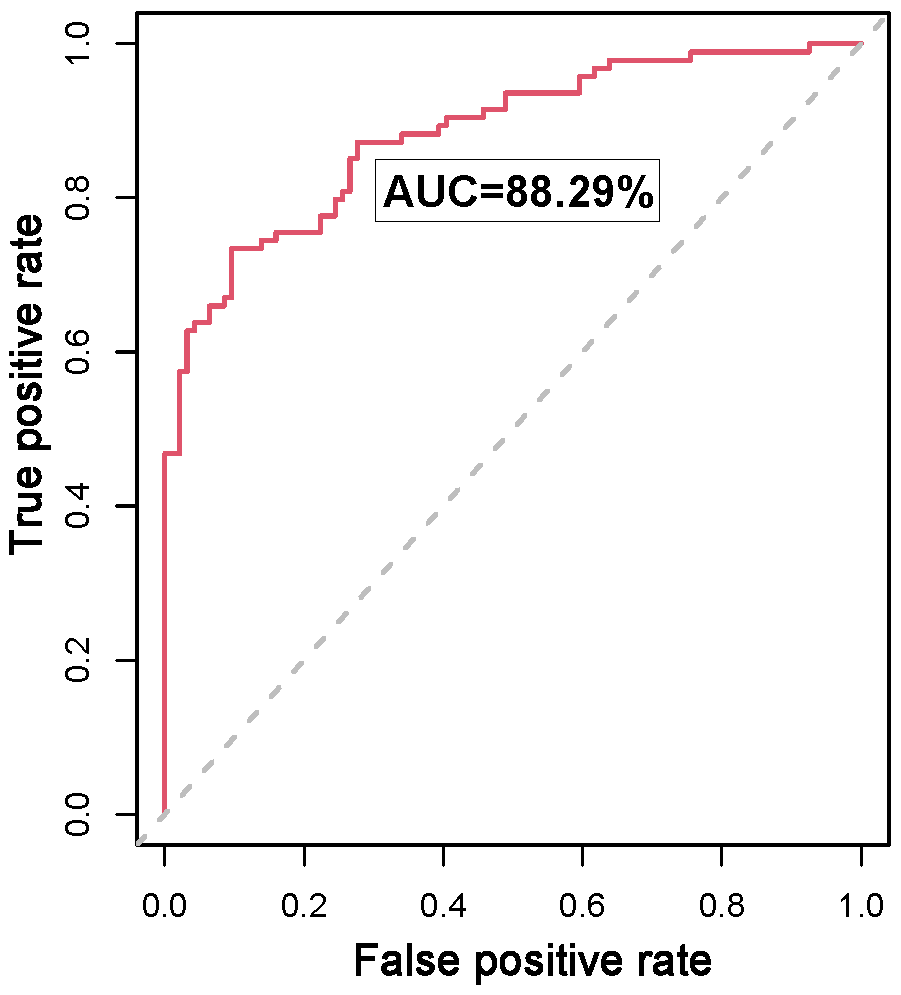

**Fig. S3.** ROC curve (red line) derived from the Random Forests model to classify *Memory* and *Control* individuals using functional connectivity, PRS, clinical and behavioral predictors. The predictive accuracy measured by the AUC was 88.29%. The diagonal line (dashed gray line) indicates the line of “no discrimination” and splits the area into the upper and lower half (50%).

#### 2.3. Top Significant Features Contributed to the Classification

The 29 significant features that were identified by the Random Forests classification and ranked based on mean minimal depth against number of decision trees are illustrated in **Fig. S4**. Minimal depth for a variable in a tree refers to the depth of the node which splits on that variable measured from the root of the tree. Lower mean minimal depth represents the higher number of observations/participants categorized into a specific group based on the variable and thus contributing to better classification accuracy, and therefore smaller depth for a feature indicates its higher importance in classification.

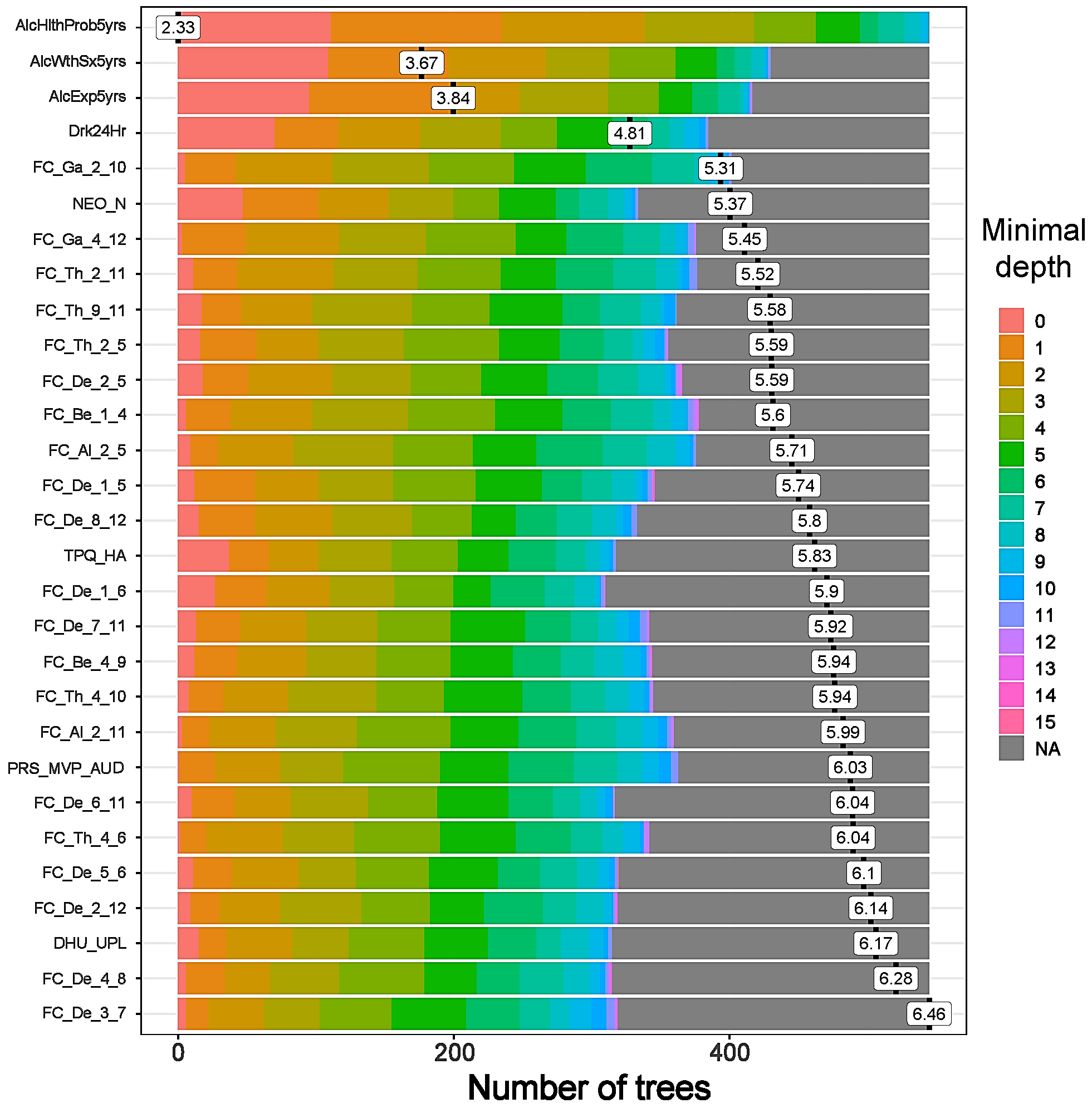

**Fig. S4.** The distribution of minimal depth among the trees of the forests for each feature is color-coded for different levels of minimal depth. The features that contributed to classifying *Memory* individuals from the *Control* participants are ranked in ascending order of minimal depth.

In order to determine the concordance of rankings between any two Random Forest (RF) parameters, correlation matrix was plotted against each parameter (see **Fig. S5)**. It was found that all of the Random Forests parameters of importance showed very high correlations among each other in ranking the features, suggesting a high concordance among these parameters in group classification.

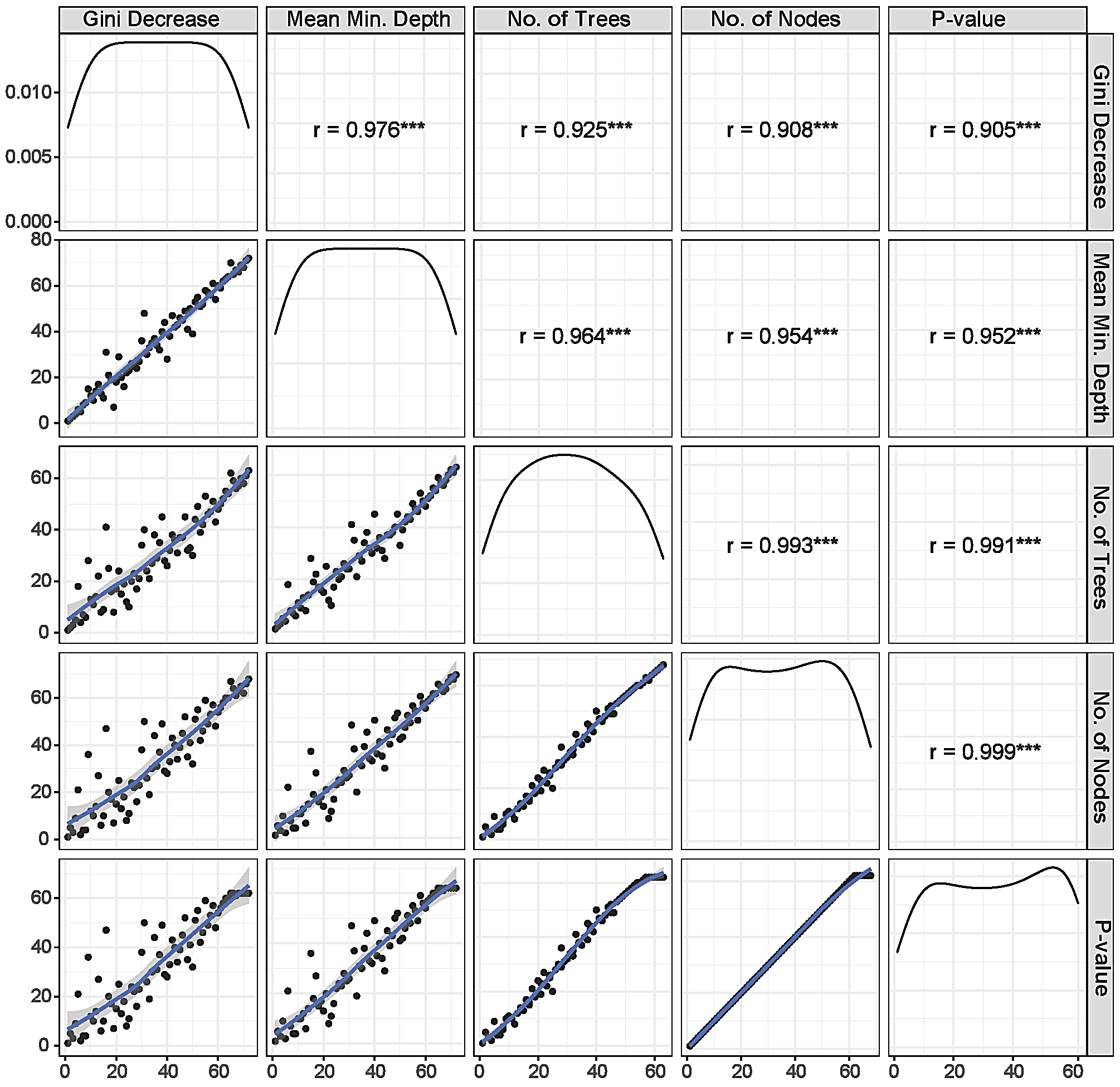

**Fig. S5:** Concordance of rankings between any two Random Forests parameters. Panels in the lower triangle of the grid show the distribution of rankings for all 72 variables (black dots) around a trend line (blue curve). The panels in the upper triangle of the grid show a correlation coefficient between the rankings of any two parameters.

### References for the Supplementary Material

[1]	Bucholz KK, Cadoret R, Cloninger CR, Dinwiddie SH, Hesselbrock VM, Nurnberger JI, Jr., Reich T, Schmidt I, Schuckit MA (1994) A new, semi-structured psychiatric interview for use in genetic linkage studies: a report on the reliability of the SSAGA. *J Stud Alcohol,* 55(2), 149-158. <https://doi.org/10.15288/jsa.1994.55.149>

[2]	Hesselbrock M, Easton C, Bucholz KK, Schuckit M, Hesselbrock V (1999) A validity study of the SSAGA--a comparison with the SCAN. *Addiction,* 94(9), 1361-1370. <https://doi.org/10.1046/j.1360-0443.1999.94913618.x>

[3]	Schuckit MA, Smith TL, Danko G, Kramer J, Bucholz KK, McCutcheon V, Chan G, Kuperman S, Hesselbrock V, Dick DM, Hesselbrock M, Porjesz B, Edenberg HJ, Nureberger JI, Jr., Gregg M, Schoen L, Kawamura M, Mendoza LA (2018) A 22-Year Follow-Up (Range 16 to 23) of Original Subjects with Baseline Alcohol Use Disorders from the Collaborative Study on Genetics of Alcoholism. *Alcohol Clin Exp Res,* 42(9), 1704-1714. <https://doi.org/10.1111/acer.13810>

[4]	Chan G, Kramer JR, Schuckit MA, Hesselbrock V, Bucholz KK, Edenberg HJ, Acion L, Langbehn D, McCutcheon V, Nurnberger JI, Jr., Hesselbrock M, Porjesz B, Bierut L, Marenna BC, Cookman A, Kuperman S (2019) A Pilot Follow-Up Study of Older Alcohol-Dependent COGA Adults. *Alcohol Clin Exp Res,* 43(8), 1759-1768. <https://doi.org/10.1111/acer.14116>

[5]	Jasper HH (1958) Report of the committee on methods of clinical examination in electroencephalography. *Electroencephalogr Clin Neurophysiol,* 10(2), 370-375. <https://doi.org/https://doi.org/10.1016/0013-4694(58)90053-1>

[6]	American Electroencephalographic Society (1994) Guideline thirteen: guidelines for standard electrode position nomenclature. American Electroencephalographic Society. *J Clin Neurophysiol,* 11(1), 111-113. <https://www.ncbi.nlm.nih.gov/pubmed/8195414>

[7]	Nuwer MR, Comi G, Emerson R, Fuglsang-Frederiksen A, Guerit JM, Hinrichs H, Ikeda A, Luccas FJ, Rappelsburger P (1998) IFCN standards for digital recording of clinical EEG. International Federation of Clinical Neurophysiology. *Electroencephalogr Clin Neurophysiol,* 106(3), 259-261. <https://www.ncbi.nlm.nih.gov/pubmed/9743285>

[8]	Begleiter H, Reich T, Hesselbrock V, Porjesz B, Li T-K, Schuckit MA, Edenberg HJ, Rice JP (1995) The Collaborative Study on the Genetics of Alcoholism. *Alcohol Health Res World,* 19(2), 228-236. <https://pubmed.ncbi.nlm.nih.gov/31798102/>

[9]	Kuperman S, Porjesz B, Arndt S, Bauer L, Begleiter H, Cizadlo T, O'Connor S, Rohrbaugh J (1995) Multi-center N400 ERP consistency using a primed and unprimed word paradigm. *Electroencephalogr Clin Neurophysiol,* 94(6), 462-470. <https://doi.org/10.1016/0013-4694(94)00312-9>

[10]	Kamarajan C, Ardekani BA, Pandey AK, Chorlian DB, Kinreich S, Pandey G, Meyers JL, Zhang J, Kuang W, Stimus AT, Porjesz B (2020) Random Forest Classification of Alcohol Use Disorder Using EEG Source Functional Connectivity, Neuropsychological Functioning, and Impulsivity Measures. *Behav Sci (Basel),* 10(3), 62. <https://doi.org/10.3390/bs10030062>

[11]	Pascual-Marqui RD (2007) Discrete, 3D distributed, linear imaging methods of electric neuronal activity. Part 1: exact, zero error localization. *arXiv,* 0710.3341. <http://arxiv.org/pdf/0710.3341>

[12]	Pascual-Marqui RD, Lehmann D, Koukkou M, Kochi K, Anderer P, Saletu B, Tanaka H, Hirata K, John ER, Prichep L, Biscay-Lirio R, Kinoshita T (2011) Assessing interactions in the brain with exact low-resolution electromagnetic tomography. *Philos Trans A Math Phys Eng Sci,* 369(1952), 3768-3784. <https://doi.org/10.1098/rsta.2011.0081>

[13]	Kamarajan C, Ardekani BA, Pandey AK, Kinreich S, Pandey G, Chorlian DB, Meyers JL, Zhang J, Bermudez E, Stimus AT, Porjesz B (2020) Random Forest Classification of Alcohol Use Disorder Using fMRI Functional Connectivity, Neuropsychological Functioning, and Impulsivity Measures. *Brain Sci,* 10(2), 115. <https://doi.org/10.3390/brainsci10020115>

[14]	Canuet L, Ishii R, Pascual-Marqui RD, Iwase M, Kurimoto R, Aoki Y, Ikeda S, Takahashi H, Nakahachi T, Takeda M (2011) Resting-state EEG source localization and functional connectivity in schizophrenia-like psychosis of epilepsy. *PLOS ONE,* 6(11), e27863. <https://doi.org/10.1371/journal.pone.0027863>

[15]	Whitfield-Gabrieli S, Nieto-Castanon A (2012) Conn: a functional connectivity toolbox for correlated and anticorrelated brain networks. *Brain Connect,* 2(3), 125-141. <https://doi.org/10.1089/brain.2012.0073>

[16]	Dansereau C, Benhajali Y, Risterucci C, Pich EM, Orban P, Arnold D, Bellec P (2017) Statistical power and prediction accuracy in multisite resting-state fMRI connectivity. *Neuroimage,* 149, 220-232. <https://doi.org/10.1016/j.neuroimage.2017.01.072>

[17]	Zuckerman M (1994) *Behavioral expressions and biosocial bases of sensation seeking.* Cambridge university press. <https://www.cambridge.org/us/academic/subjects/life-sciences/animal-behaviour/behavioral-expressions-and-biosocial-bases-sensation-seeking?format=HB&isbn=9780521432009>

[18]	Cloninger CR, Przybeck TR, Svrakic DM, Wetzel RD (1994) *The Temperament and Character Inventory (TCI): A Guide to Its Development and Use.* Washington University, St. Louis, MO. <https://www.researchgate.net/publication/264329741_TCI-Guide_to_Its_Development_and_Use>

[19]	DeLongis A, Folkman S, Lazarus RS (1988) The impact of daily stress on health and mood: psychological and social resources as mediators. *J Pers Soc Psychol,* 54(3), 486-495. <https://doi.org/10.1037//0022-3514.54.3.486>

[20]	Costa PT, Jr., McCrae RR (1986) Cross-sectional studies of personality in a national sample: 1. Development and validation of survey measures. *Psychol Aging,* 1(2), 140-143. <https://www.ncbi.nlm.nih.gov/pubmed/3267390>

[21]	Procidano ME, Heller K (1983) Measures of perceived social support from friends and from family: three validation studies. *Am J Community Psychol,* 11(1), 1-24. <https://doi.org/10.1007/BF00898416>

[22]	Brown SA, Christiansen BA, Goldman MS (1987) The Alcohol Expectancy Questionnaire: an instrument for the assessment of adolescent and adult alcohol expectancies. *J Stud Alcohol,* 48(5), 483-491. <https://doi.org/10.15288/jsa.1987.48.483>

[23]	Schuckit MA, Tipp JE, Smith TL, Wiesbeck GA, Kalmijn J (1997) The relationship between Self-Rating of the Effects of alcohol and alcohol challenge results in ninety-eight young men. *J Stud Alcohol,* 58(4), 397-404. <https://doi.org/10.15288/jsa.1997.58.397>

[24]	Edenberg HJ, Koller DL, Xuei X, Wetherill L, McClintick JN, Almasy L, Bierut LJ, Bucholz KK, Goate A, Aliev F, Dick D, Hesselbrock V, Hinrichs A, Kramer J, Kuperman S, Nurnberger JI, Jr., Rice JP, Schuckit MA, Taylor R, Todd Webb B, Tischfield JA, Porjesz B, Foroud T (2010) Genome-wide association study of alcohol dependence implicates a region on chromosome 11. *Alcohol Clin Exp Res,* 34(5), 840-852. <https://doi.org/10.1111/j.1530-0277.2010.01156.x>

[25]	Wang JC, Foroud T, Hinrichs AL, Le NX, Bertelsen S, Budde JP, Harari O, Koller DL, Wetherill L, Agrawal A, Almasy L, Brooks AI, Bucholz K, Dick D, Hesselbrock V, Johnson EO, Kang S, Kapoor M, Kramer J, Kuperman S, Madden PA, Manz N, Martin NG, McClintick JN, Montgomery GW, Nurnberger JI, Jr., Rangaswamy M, Rice J, Schuckit M, Tischfield JA, Whitfield JB, Xuei X, Porjesz B, Heath AC, Edenberg HJ, Bierut LJ, Goate AM (2013) A genome-wide association study of alcohol-dependence symptom counts in extended pedigrees identifies C15orf53. *Mol Psychiatry,* 18(11), 1218-1224. <https://doi.org/10.1038/mp.2012.143>

[26]	Baurley JW, Edlund CK, Pardamean CI, Conti DV, Bergen AW (2016) Smokescreen: a targeted genotyping array for addiction research. *BMC Genomics,* 17, 145. <https://doi.org/10.1186/s12864-016-2495-7>

[27]	Delaneau O, Howie B, Cox AJ, Zagury JF, Marchini J (2013) Haplotype estimation using sequencing reads. *Am J Hum Genet,* 93(4), 687-696. <https://doi.org/10.1016/j.ajhg.2013.09.002>

[28]	Das S, Forer L, Schonherr S, Sidore C, Locke AE, Kwong A, Vrieze SI, Chew EY, Levy S, McGue M, Schlessinger D, Stambolian D, Loh PR, Iacono WG, Swaroop A, Scott LJ, Cucca F, Kronenberg F, Boehnke M, Abecasis GR, Fuchsberger C (2016) Next-generation genotype imputation service and methods. *Nat Genet,* 48(10), 1284-1287. <https://doi.org/10.1038/ng.3656>

[29]	Meyers JL, Zhang J, Wang JC, Su J, Kuo SI, Kapoor M, Wetherill L, Bertelsen S, Lai D, Salvatore JE, Kamarajan C, Chorlian D, Agrawal A, Almasy L, Bauer L, Bucholz KK, Chan G, Hesselbrock V, Koganti L, Kramer J, Kuperman S, Manz N, Pandey A, Seay M, Scott D, Taylor RE, Dick DM, Edenberg HJ, Goate A, Foroud T, Porjesz B (2017) An endophenotype approach to the genetics of alcohol dependence: a genome wide association study of fast beta EEG in families of African ancestry. *Mol Psychiatry,* 22(12), 1767-1775. <https://doi.org/10.1038/mp.2016.239>

[30]	Wetherill L, Agrawal A, Kapoor M, Bertelsen S, Bierut LJ, Brooks A, Dick D, Hesselbrock M, Hesselbrock V, Koller DL, Le N, Nurnberger JI, Jr., Salvatore JE, Schuckit M, Tischfield JA, Wang JC, Xuei X, Edenberg HJ, Porjesz B, Bucholz K, Goate AM, Foroud T (2015) Association of substance dependence phenotypes in the COGA sample. *Addict Biol,* 20(3), 617-627. <https://doi.org/10.1111/adb.12153>

[31]	Kranzler HR, Zhou H, Kember RL, Vickers Smith R, Justice AC, Damrauer S, Tsao PS, Klarin D, Baras A, Reid J, Overton J, Rader DJ, Cheng Z, Tate JP, Becker WC, Concato J, Xu K, Polimanti R, Zhao H, Gelernter J (2019) Genome-wide association study of alcohol consumption and use disorder in 274,424 individuals from multiple populations. *Nat Commun,* 10(1), 1499. <https://doi.org/10.1038/s41467-019-09480-8>

[32]	Gelernter J, Sun N, Polimanti R, Pietrzak RH, Levey DF, Lu Q, Hu Y, Li B, Radhakrishnan K, Aslan M, Cheung KH, Li Y, Rajeevan N, Sayward F, Harrington K, Chen Q, Cho K, Honerlaw J, Pyarajan S, Lencz T, Quaden R, Shi Y, Hunter-Zinck H, Gaziano JM, Kranzler HR, Concato J, Zhao H, Stein MB, Department of Veterans Affairs Cooperative Studies P, Million Veteran P (2019) Genome-wide Association Study of Maximum Habitual Alcohol Intake in >140,000 U.S. European and African American Veterans Yields Novel Risk Loci. *Biol Psychiatry,* 86(5), 365-376. <https://doi.org/10.1016/j.biopsych.2019.03.984>

[33]	Walters RK, Polimanti R, Johnson EC, McClintick JN, Adams MJ, Adkins AE, Aliev F, Bacanu SA, Batzler A, Bertelsen S, Biernacka JM, Bigdeli TB, Chen LS, Clarke TK, Chou YL, Degenhardt F, Docherty AR, Edwards AC, Fontanillas P, Foo JC, Fox L, Frank J, Giegling I, Gordon S, Hack LM, Hartmann AM, Hartz SM, Heilmann-Heimbach S, Herms S, Hodgkinson C, Hoffmann P, Jan Hottenga J, Kennedy MA, Alanne-Kinnunen M, Konte B, Lahti J, Lahti-Pulkkinen M, Lai D, Ligthart L, Loukola A, Maher BS, Mbarek H, McIntosh AM, McQueen MB, Meyers JL, Milaneschi Y, Palviainen T, Pearson JF, Peterson RE, Ripatti S, Ryu E, Saccone NL, Salvatore JE, Sanchez-Roige S, Schwandt M, Sherva R, Streit F, Strohmaier J, Thomas N, Wang JC, Webb BT, Wedow R, Wetherill L, Wills AG, andMe Research T, Boardman JD, Chen D, Choi DS, Copeland WE, Culverhouse RC, Dahmen N, Degenhardt L, Domingue BW, Elson SL, Frye MA, Gabel W, Hayward C, Ising M, Keyes M, Kiefer F, Kramer J, Kuperman S, Lucae S, Lynskey MT, Maier W, Mann K, Mannisto S, Muller-Myhsok B, Murray AD, Nurnberger JI, Palotie A, Preuss U, Raikkonen K, Reynolds MD, Ridinger M, Scherbaum N, Schuckit MA, Soyka M, Treutlein J, Witt S, Wodarz N, Zill P, Adkins DE, Boden JM, Boomsma DI, Bierut LJ, Brown SA, Bucholz KK, Cichon S, Costello EJ, de Wit H, Diazgranados N, Dick DM, Eriksson JG, Farrer LA, Foroud TM, Gillespie NA, Goate AM, Goldman D, Grucza RA, Hancock DB, Harris KM, Heath AC, Hesselbrock V, Hewitt JK, Hopfer CJ, Horwood J, Iacono W, Johnson EO, Kaprio JA, Karpyak VM, Kendler KS, Kranzler HR, Krauter K, Lichtenstein P, Lind PA, McGue M, MacKillop J, Madden PAF, Maes HH, Magnusson P, Martin NG, Medland SE, Montgomery GW, Nelson EC, Nothen MM, Palmer AA, Pedersen NL, Penninx B, Porjesz B, Rice JP, Rietschel M, Riley BP, Rose R, Rujescu D, Shen PH, Silberg J, Stallings MC, Tarter RE, Vanyukov MM, Vrieze S, Wall TL, Whitfield JB, Zhao H, Neale BM, Gelernter J, Edenberg HJ, Agrawal A (2018) Transancestral GWAS of alcohol dependence reveals common genetic underpinnings with psychiatric disorders. *Nat Neurosci,* 21(12), 1656-1669. <https://doi.org/10.1038/s41593-018-0275-1>

[34]	Ge T, Chen CY, Ni Y, Feng YA, Smoller JW (2019) Polygenic prediction via Bayesian regression and continuous shrinkage priors. *Nat Commun,* 10(1), 1776. <https://doi.org/10.1038/s41467-019-09718-5>

[35]	Ruan Y, Lin Y-F, Feng Y-CA, Chen C-Y, Lam M, Guo Z, He L, Sawa A, Martin AR, Qin S, Huang H, Ge T (2021) Improving Polygenic Prediction in Ancestrally Diverse Populations. *medRxiv*, 2020.2012.2027.20248738. <https://doi.org/10.1101/2020.12.27.20248738>

[36]	Lai D, Johnson EC, Colbert S, Pandey G, Chan G, Bauer L, Francis MW, Hesselbrock V, Kamarajan C, Kramer J, Kuang W, Kuo S, Kuperman S, Liu Y, McCutcheon V, Pang Z, Plawecki MH, Schuckit M, Tischfield J, Wetherill L, Zang Y, Edenberg HJ, Porjesz B, Agrawal A, Foroud T (2022) Evaluating risk for alcohol use disorder: Polygenic risk scores and family history. *Alcohol Clin Exp Res,* 46(3), 374-383. <https://doi.org/10.1111/acer.14772>

[37]	Lai D, Schwantes-An TH, Abreu M, Chan G, Hesselbrock V, Kamarajan C, Liu Y, Meyers JL, Nurnberger JI, Jr., Plawecki MH, Wetherill L, Schuckit M, Zhang P, Edenberg HJ, Porjesz B, Agrawal A, Foroud T (2022) Gene-based polygenic risk scores analysis of alcohol use disorder in African Americans. *Transl Psychiatry,* 12(1), 266. <https://doi.org/10.1038/s41398-022-02029-2>

[38]	Nguyen C, Wang Y, Nguyen HN (2013) Random forest classifier combined with feature selection for breast cancer diagnosis and prognostic. *J Biomed Sci Eng,* 06(05), 551-560. <https://doi.org/10.4236/jbise.2013.65070>

[39]	Chandrashekar G, Sahin F (2014) A survey on feature selection methods. *Computers & Electrical Engineering,* 40(1), 16-28. <https://doi.org/10.1016/j.compeleceng.2013.11.024>

[40]	Cai J, Luo JW, Wang SL, Yang S (2018) Feature selection in machine learning: A new perspective. *Neurocomputing,* 300, 70-79. <https://doi.org/10.1016/j.neucom.2017.11.077>

[41]	Friedman J, Hastie T, Tibshirani R (2010) Regularization Paths for Generalized Linear Models via Coordinate Descent. *J Stat Softw,* 33(1), 1-22. <https://www.ncbi.nlm.nih.gov/pubmed/20808728>

[42]	Fonti V, Belitser E (2017) Feature selection using LASSO. Accessed on 2019-June-01. Accessed at: <https://www.researchgate.net/profile/David-Booth-7/post/Regression-of-pairwise-trait-similarity-on-similarity-in-personal-attributes/attachment/5b18368d4cde260d15e3a4e3/AS%3A634606906785793%401528313485788/download/werkstuk-fonti_tcm235-836234.pdf>

[43]	Breiman L (2001) Random forests. *Machine learning,* 45(1), 5-32. <https://doi.org/10.1023/A:1010933404324>

[44]	Qi Y (2012) Random Forest for Bioinformatics. In: Zhang C, Ma Y (eds.): *Ensemble Machine Learning*, Springer US, Boston, MA, pp. 307-323. <https://doi.org/https://doi.org/10.1007/978-1-4419-9326-7_11>

[45]	Couronne R, Probst P, Boulesteix AL (2018) Random forest versus logistic regression: a large-scale benchmark experiment. *BMC Bioinformatics,* 19(1), 270. <https://doi.org/10.1186/s12859-018-2264-5>

[46]	Strobl C, Malley J, Tutz G (2009) An introduction to recursive partitioning: Rationale, application, and characteristics of classification and regression trees, bagging, and random forests. *Psychol Methods,* 14(4), 323-348. <https://doi.org/10.1037/a0016973>

[47]	Breiman L, Cutler A (2019) *Random Forest*. Accessed on 2019-June-01. Webpage: <https://www.stat.berkeley.edu/~breiman/RandomForests/cc_home.htm#ooberr>
